# Beyond Bisulfite Sequencing: Resolving 5-hmC with Nanopore Sequencing Unmasks the True Methylation Entropy Landscape

**DOI:** 10.64898/2026.07.08.736699

**Authors:** Uri Bertocchi, Eyal Katz, Jonathan Jeffet, Assaf Grunwald, Neiv Gabay, Jasline Deek, Sujal Verma, Amit Shwartz, Gali Umschweif-Nevo, Bernard Lerer, Yael Roichman, Yuval Ebenstein

**Affiliations:** Sagol School of Neuroscience, Tel Aviv University, 55 Chaim Levanon St, Tel Aviv-Yafo 6997801, Israel; Raymond & Beverly Sackler School of Physics & Astronomy, Faculty of Exact Sciences, Tel Aviv University, 55 Chaim Levanon St, Tel Aviv-Yafo 6997801, Israel; Raymond & Beverly Sackler School of Chemistry, Faculty of Exact Sciences, Tel Aviv University, 55 Chaim Levanon St, Tel Aviv-Yafo 6997801, Israel; School of Biomedical Engineering, The Iby and Aladar Fleischman Faculty of Engineering, Tel Aviv University, 55 Chaim Levanon St, Tel Aviv-Yafo 6997801, Israel; School of Pharmacy, Faculty of Medicine, The Hebrew University of Jerusalem, Ein Kerem Campus, Kalman Ya’akov Man St, Jerusalem 9112102, Israel; Hadassah BrainLabs Center for Psychedelic Research, Hadassah–Hebrew University Medical Center, Ein Kerem Campus, Kalman Ya’akov Man St, Jerusalem 91120, Israel

**Keywords:** DNA methylation, 5-hydroxymethylcytosine, nanopore sequencing, methylation entropy, psilocybin, clear cell renal cell carcinoma

## Abstract

DNA methylation dynamically regulates cellular function and phenotype. At the tissue level, stochasticity in methylation patterns, as measured by methylation entropy, drives plasticity, development, cancer, and aging. Modulation of methylation patterns is facilitated by erasure of 5-methylcytosine (5mC) via the oxidized intermediate 5-hydroxymethylcytosine (5hmC). Bisulfite sequencing cannot distinguish the two modifications, labeling them both as 5-mC. We quantitatively analyze the effect of this historical conflation on the genome-wide distribution of methylation levels and methylation entropy. Using nanopore sequencing with direct 5mC and 5hmC calling, we compare True-mC to bisulfite-like analysis in two model systems, kidney cancer and the mouse medial prefrontal cortex. We show that bisulfite sequencing introduces systematic, tissue-specific shifts in methylation distribution that affect the mechanistic interpretation of the underlying biology. The distortion scaled with endogenous 5hmC content: substantial in brain, where over half of the entropy-associated gene ontology terms recovered under bisulfite-like analysis were absent under True-mC, and minimal in the low-5hmC kidney cancer sample, which delimits the regime in which bisulfite-derived entropy remains interpretable. Together, these findings establish that True-5mC-based methylation entropy redefines the physical mapping of certain epigenomes, demonstrating that in some contexts, what has previously been interpreted as stochastic maintenance failure is frequently the structured signature of distinct and mechanistically interpretable cytosine biochemistry.

## INTRODUCTION

The epigenome serves as the critical interface between static genetic information and dynamic cellular phenotypes. In classical models of development, such as Waddington’s epigenetic landscape(1), cellular differentiation is driven by the progressive restriction of epigenetic plasticity. While stable DNA methylation level patterns define cell identity, it is increasingly recognized that methylation stochasticity, or “noise”, is a key driver of cellular plasticity in both normal development and disease pathogenesis(2–5). Principles from information theory and statistical physics have long been applied to whole-genome methylation data to rigorously quantify methylation stochasticity(6, 7). By calculating methylation entropy (ME), researchers can construct epigenetic energy landscapes, in which high entropy reflects a highly plastic or disordered state, and low entropy indicates a rigidly stabilized, highly correlated genomic architecture(8, 9).

The empirical calculation of global ME has historically relied on bisulfite sequencing, introducing a fundamental chemical blind spot. Standard bisulfite conversion cannot distinguish between 5-methylcytosine (5mC) and its oxidized derivative, 5-hydroxymethylcytosine (5hmC), interpreting both modifications indistinguishably as a protected cytosine(10, 11). Unlike 5mC, which encodes stable epigenetic memory through *de-novo* and maintenance methylation, 5hmC serves a dual biological role: it functions both as a stable epigenetic mark with dedicated protein binders and readers(12, 13), and as an obligate intermediate in TET-mediated active demethylation(14, 15), with the relative contribution of each role being tissue-specific(16, 17). Because 5hmC is deposited in a coordinated, enzymatically directed manner(18, 19), pooling it with the inherently more heterogeneous 5mC maintenance signal can introduce a systematic, structured bias rather than random noise, skewing the empirical distribution of methylation entropy versus mean methylation level (MML). This is amplified in tissues characterized by high endogenous 5hmC levels, such as the mammalian brain (20, 21), where massive epigenomic reconfiguration drives neuroplasticity (22).

To systematically resolve this artifact, we used Oxford Nanopore Technologies (ONT) long-read sequencing, which detects nucleotide-specific molecular signatures from native DNA and distinguishes naturally occurring cytosine modifications, such as 5mC(23, 24) and 5hmC(25–28), on the same molecule. Moreover, long-read sequencing preserves the complete methylation pattern of each molecule(23). This long-range single-base resolution allows distinct epialleles(29, 30) to be distinguished within a heterogeneous cell population(7, 30), as well as lineage tracing in epigenetic inheritance studies(31).

The reference implementation for information-theoretic methylation analysis is informME, which fits a one-dimensional Ising model to WGBS reads within genomic windows and reports the mean methylation level (MML) and the normalized methylation entropy for each analysis region(8, 32), with differential analysis performed on the Jensen-Shannon distance between the corresponding methylation-level distributions. The MML-ME representation used here is therefore not itself new; what is new is the state space over which it is computed. Because informME and its derivatives operate on bisulfite reads, 5mC and 5hmC enter the model as a single protected state, so the entropy is defined over a chemically ambiguous binary alphabet. Existing nanopore tools (Modkit(33, 34), PoreMeth2(35)) read native modification calls but were not designed to isolate the 5hmC contribution and do not implement a Shannon-entropy formulation over resolved cytosine states. We therefore developed ShannonPore, which computes the MML-ME phase space from true 5mC calls while retaining 5hmC as a separate state, and which additionally reports the bisulfite-like projection of the same reads. This allows the two chemistries to be contrasted on identical molecules rather than across separate experiments and sample preparations, a comparison that cannot be constructed from bisulfite data alone (Supplementary Fig. 1). We applied our analysis to two biological paradigms selected for their contrasting 5hmC dynamics. First, we analyzed a clear cell renal cell carcinoma (ccRCC) sample with low-5hmC(36, 37), and driven by deterministic chromosomal deletions(38–40). One validated deletion involves the loss of SETD2, which affects DNA methyltransferase activity and results in biologically anchored redistribution of methylation(41, 42). Second, we evaluated psilocybin-induced neuroplasticity in the mouse prefrontal cortex(22, 43, 44). This tissue represents an environment with massive endogenous 5hmC flux(20, 21, 37), presenting the most acute challenge for bisulfite conflation.

## MATERIALS AND METHODS

Two datasets were analyzed: matched tumor and normal kidney tissue from a ccRCC patient, and medial prefrontal cortex tissue from psilocybin-treated or saline-treated mice (n=5 each). Both were processed through the same nanopore sequencing and methylation entropy analysis pipeline.

### ccRCC Clinical Samples

#### Patient and Tissue

The tumor/normal pair was acquired and sequenced in a previous project in accordance with the Declaration of Helsinki(45). In short, matched tumor and normal adjacent tissue were obtained from an 82-year-old male patient undergoing radical nephrectomy. The tumor was diagnosed histologically as ccRCC, eosinophilic variant, pT3a stage. Tissues were stored at −80°C from the time of surgery until analysis. Institutional review boards approved sample collection and handling, and DNA extraction.

Ultra-high-molecular-weight DNA was extracted from fresh-frozen tissue using the SP Tissue and Tumor DNA Isolation Kit (Bionano Genomics) according to the manufacturer’s protocol. The genomic architecture of these samples was rigorously characterized using integrated Optical Genome Mapping and Nanopore-derived SNV analysis, with results previously validated and published(46, 47); see Supplementary Fig. 13 for 3p loss.

### Psilocybin Mouse Model

#### Animals

Five male C57BL/6J mice per experimental group (*n* = 5 control; *n* = 5 psilocybin-treated) aged 11 weeks were used. All animal procedures were approved by the institutional animal care and use committee of the Hebrew University (AAALAC accreditation #1285; Ethics approval number MD-23-17322-4). Animal studies were reported in accordance with the National Institutes of Health Guide for the Care and Use of Laboratory Animals guidelines(48). The animals were housed in a temperature-controlled environment (25D±D2D°C, relative humidity 55D±D15%) with a 12-h light/dark cycle. All mice were fed a standard laboratory diet of nutrient-rich pellets ad libitum.

#### Psilocybin Administration

Psilocybin (Usona Institute, LOT AM50167; MW 284.25; purity 98.75%) was dissolved in 0.9% NaCl (saline) vehicle at 0.44 mg/mL and administered intraperitoneally (IP) at 4.4 mg/kg. This dose is within the therapeutic range used in clinical trials for treatment-resistant depression, and was selected to achieve a clinically translatable neuroplastic stimulus(49, 50). The injection volume was standardized to 10 µL per gram body weight to account for inter-animal variation in mass. Control animals received an equivalent volume of saline by the same route and under the same handling conditions.

#### Tissue Collection

Animals were euthanized 72 hours post-injection. This time point was chosen because the acute serotonergic response resolves within hours of administration, and prior transcriptomic studies show that gene expression changes associated with synaptic remodeling stabilize within this window, making it appropriate for capturing the consolidation of epigenetic changes(22, 43, 51, 52). The mPFC was dissected and put immediately in liquid nitrogen to preserve chromatin and RNA integrity. Five animals were analyzed per group. Although group sizes were small, the high-coverage single-molecule nature of nanopore sequencing generates average per-CpG read depths of 23.5× (see Supplementary Table 1).

#### DNA Extraction

Brain tissue was homogenized using ZR BashingBead Lysis Tubes (2 mm), and genomic DNA was extracted using the Zymo Quick-DNA/RNA Miniprep Plus Kit according to the manufacturer’s protocol.

### Nanopore Sequencing

Libraries of ccRCC and mouse brain samples were prepared using the Ligation Sequencing Kit V14 (SQK-LSK114, Oxford Nanopore Technologies) and sequenced on a PromethION device with R10.4.1 flow cells.

### Computational Pipeline

#### Basecalling and Modification Detection

Reads were basecalled using the Dorado basecalling model (dna_r10.4.1_e8.2_400bps_sup@v5.2.0@5mCG_5hmCG@v2), which simultaneously calls 5mC and 5hmC at CpG dinucleotides. The superhigh-accuracy model was used to maximize the fidelity of modification detection. Because nanopore 5hmC-calling accuracy is model-version-dependent, the per-call accuracy reported here corresponds specifically to the R10.4.1/Dorado v5.2.0 5mCG_5hmCG model. Reads were aligned to the human reference genome (hg38) for the ccRCC dataset and to the mouse reference genome (mm10) for the psilocybin dataset using Dorado’s in-house minimap calling. Per-read CpG modification probabilities were extracted from aligned BAM files using modkit extract (v0.6.1), producing per-read TSV files retaining the identity of each cytosine modification call at single-molecule resolution.

#### Methylation Entropy and Mean Methylation Level

MML and ME were computed using ShannonPore (v0.1.0; https://github.com/ebensteinLab/ShannonPore). For the ternary entropy comparison between bisulfite-like, true-mC, and ternary frameworks, bins of 3 consecutive CpGs were used with a minimum coverage threshold of 27×. This threshold ensures that all 3³ = 27 possible ternary read patterns can, in principle, be observed in each bin, preventing underestimation of entropy due to incomplete pattern sampling. For all other analyses, bins of 4 consecutive CpGs were used with a minimum coverage of 16×, the threshold at which all 2D = 16 binary read patterns can be observed. CpGs were defined per dinucleotide on the reference; methylation calls from minus-strand reads were shifted by −1 to align with the plus-strand C, so each CpG locus is represented once and receives one observation per read with coverage pooled across both strands. After all quality filters, 4,376,236 4-CpG bins were retained for the kidney dataset and 3,591,369 for the psilocybin mouse dataset (requiring coverage in at least 3 of 5 samples per group).

Bisulfite-like entropy was computed by converting all nanopore 5hmC calls to 5mC before entropy calculation, emulating the chemical indistinguishability of 5mC and 5hmC under bisulfite treatment. To do this, ShannonPore leverages the ONT modkit modbam functionality, specifically using the “adjust-mods” and “convert” commands to reassign 5hmC (the “h” code) to 5mC (the “m” code). True-mC entropy was computed by converting all 5hmC calls to unmodified C. Ternary entropy retained 5mC, 5hmC, and unmodified C as three distinct states. All three conversion schemes are implemented within ShannonPore and available in open source.

#### Gene Ontology Enrichment Analysis

GO enrichment analysis was performed using gProfiler2(53) with genomic bin coordinates supplied directly as input, enabling region-based pathway recovery without requiring prior gene assignment. This approach captures regulatory signals from bins that overlap gene bodies, promoters, and intergenic elements without imposing a gene-centric selection step. Significance was assessed using a false discovery rate threshold of *q* < .05.

## RESULTS

DNA methylation and hydroxymethylation occur predominantly at cytosines that are followed by guanine (CpG dinucleotides) embedded within the genetic sequence (54, 55). The most chemically complete representation of the system is a ternary model, in which C, 5mC, and 5hmC are each treated as distinct states. Bisulfite conversion does not distinguish between the two cytosine modifications, marking them both as methylated. For the sake of measuring ME and MML, we consider a chain of methylation sites that can take one of three forms: C, 5mC, and 5hmC. Each position may be assigned an MML by averaging the methylation state across all sequencing reads to a value between 0 and 1 (often called the beta value). Beyond the MML, additional information content lies in the methylation pattern. The degree of stochasticity in the pattern of a certain region may be calculated by the Shannon entropy formula (see Figure 1a for an illustration; refer to Supplementary Note 1 for a detailed explanation):

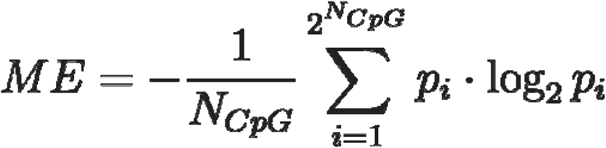

where is the number of methylation sites within the sequence over which the ME is calculated, and is the probability of a specific methylation pattern within that sequence (Fig. 1a).

**Fig. 1:**
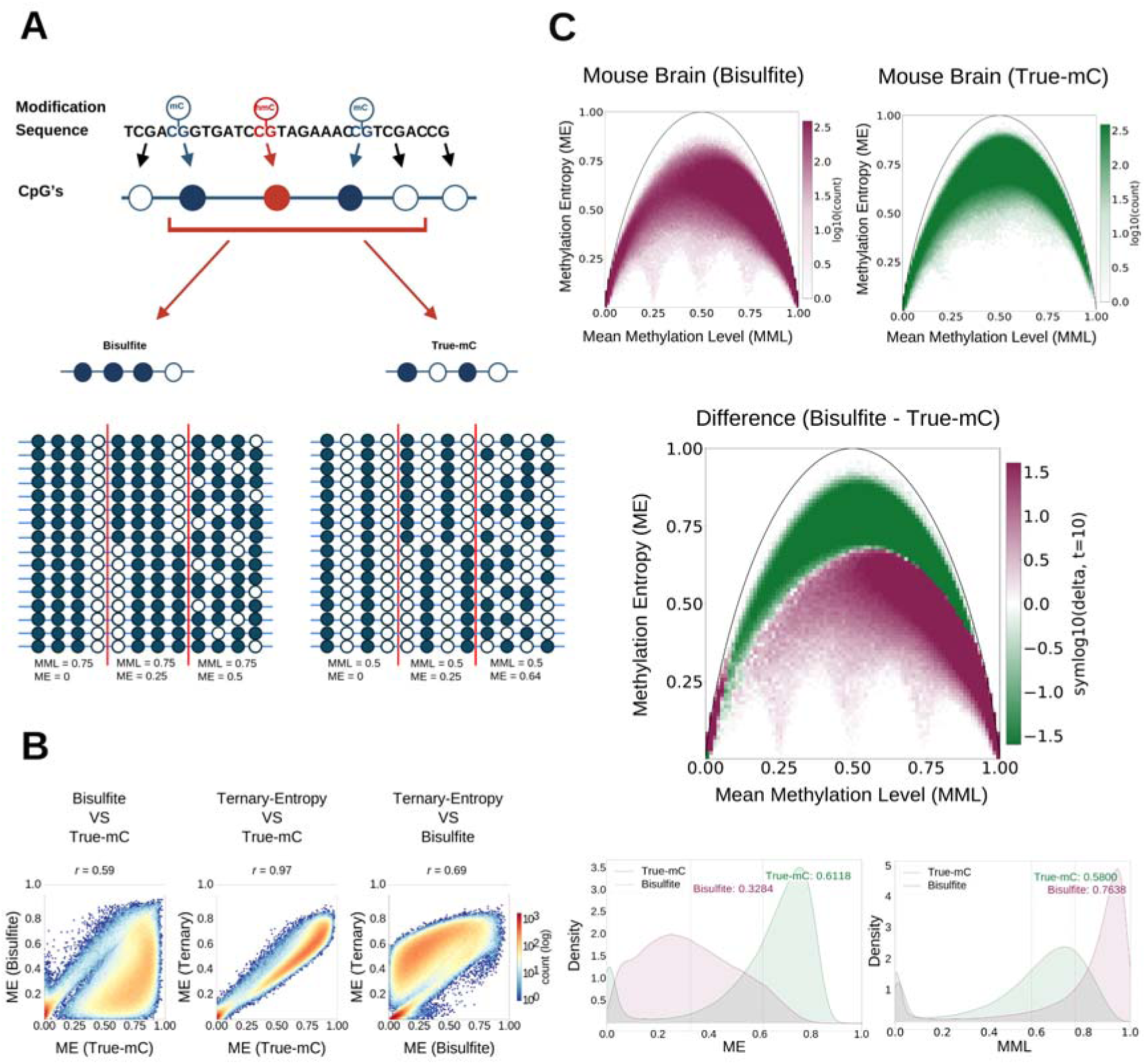
Chemical resolution of cytosine modifications and their effect on methylation entropy. **a** schematic of a 4-CpG locus carrying one 5hmC alongside 5mC sites. Bisulfite sequencing marks both modifications as methylated, resulting in higher apparent MML and lower apparent ME relative to true-mC measurements. **b** Correlation between three parallel ME frameworks computed from the same nanopore data on 3-CpG windows (mouse mPFC; N = 5, minimum coverage 27×). True-mC: 5hmC treated as unmodified C. Bisulfite-like: 5hmC treated as 5mC to emulate bisulfite chemistry. Ternary: 5mC, 5hmC, and unmodified C are each retained as distinct states. **c** Phase-space distributions (ME vs. MML, 4-CpG bins) in brain mPFC under bisulfite-like (purple) and true-mC (green). Top: 2D Phase-space maps of bisulfite-like and true-mC. Middle: signed log_₁₀_ difference map (bisulfite-like minus true-mC); purple indicates bins more represented in the bisulfite data, green indicates bins more abundant in the true-mC data. Bottom: density distribution of true-mC (purple) vs bisulfite-like (green) in MML (left plot), and ME (right plot).

We first consider the mathematical and analytical framework for methylation entropy analysis. In the context of methylation, Shannon entropy measures the diversity of methylation patterns across multiple sites within the same genomic unit. Ideally, one would like to have ME calculated for biologically distinct regions along the genome, such as regulatory elements spanning many methylation sites.

However, calculating ME across many methylation sites yields numerous possible configurations that are impossible to reliably sample with conventional sequencing depth. Consequently, mapping ME requires partitioning the genomic sequence into discrete windows, or bins, containing a specific CpG count. Historically, two main binning strategies have been used for this purpose. The first divides the genome into fixed-length windows, such as 150-bp intervals(8, 32, 56); the second divides the genome into windows containing a fixed number of CpGs (6, 57). To determine the optimal binning strategy for our nanopore datasets, we performed error simulations using the hg38 reference genome sequence at 30x sequencing coverage (Supplementary Figs. 2-5). Fixed-length genomic windows introduced an error that depends on CpG density. CpG-rich regions contained many CpGs per bin and therefore required unrealistically high coverage to sample the full entropy space. At the same time, CpG-poor regions contained too few CpGs to provide stable entropy estimates. The simulations show that at constant CpGs per bin the lowest error in ME is achieved for 3 CpG bins. However, 3 CpG bins provide only 2^3^=8 methylation configurations and capture shorter range correlations. Consequently, a calculation that accounts for estimation error shows that the usable information content in 4 CpG bins is higher despite the larger estimation error (Supplementary Fig. 4). We therefore propose 4-CpG bins as a practical compromise between entropy-estimation accuracy, accessible ME levels, and realistic nanopore sequencing depth. This configuration preserves sufficient pattern complexity while remaining compatible with the typical 20–30× coverage achieved in most common sequencing experiments.

The most complete framework for 5hmC-aware methylation entropy is a ternary model in which each CpG site can be in one of three states: C, 5mC, or 5hmC. However, a three-state 4-CpG bin yields 3D = 81 possible patterns, which requires unrealistically high sequencing coverage. While bisulfite sequencing reduces this model to two states by combining 5mC and 5hmC, an alternative approach, enabled by nanopore sequencing, is to consider only true 5mC (True-mC), combining 5hmC with unmodified C.

This raises the question of which of the two binary approaches, that do not explicitly count 5hmC, more closely represents the underlying biology. Given that the effect of bisulfite-based analysis is enhanced in high-5hmC tissues, such as the brain, where a large fraction of modified cytosines is 5hmC(20, 21, 37), we assessed concordance with the ternary model using ONT sequencing of five mouse medial prefrontal cortices (mPFC). By calculating genome-wide entropy across 3-CpG bins, we reduced the ternary pattern space to 3³ = 27 possible configurations and used high coverage genomic regions for analysis. Figure 1b presents pairwise correlations among the three entropy calculations for these samples, with True-mC best projecting the ternary model, with an r = 0.94, compared to r = 0.71 for bisulfite. This indicates that when 5hmC content is high, True-mC better preserves the entropy structure obtained when 5hmC is explicitly modeled; see Supplementary Fig. 6 for the cytosine modification composition across the study’s samples and Supplementary Fig. 7 for the prevalence and ternary phased-space plots of C, 5mC, and 5hmC in the samples.

To visualize the full methylation information, we represent methylation bins in a smooth two-dimensional phase space (Figure 1c), with each bin shown by its calculated MML on the x-axis and its ME on the y-axis. The black arc denotes the maximum theoretical Shannon entropy for a given MML obtained for completely random methylation. The extremes of the phase space are locked at fully methylated or unmethylated, which have zero entropy, and the half-methylated configuration has the highest number of possible methylation patterns. Looking at the collapsed distributions (lower right panel of Fig 1c), the MML distribution peak shifts towards lower methylation for True-mC, as expected from omitting highly hydroxymethylated cytosines from the bisulfite-methylated pool. ME shows a more interesting change, where a broad, unimodal distribution leaning towards intermediate-low entropy for bisulfite shifts to a bimodal distribution weighted towards high-entropy bins for True-mC. The absolute phase-space difference map (bisulfite-like minus True-mC) explicitly defines the geometric direction of the effect: in bisulfite, excess bins accumulate in the high-MML, low-ME corner of phase space and in the mid-low ME region. Such a profile would conventionally be interpreted as rigidly maintained and highly correlated methylation. True-mC reveals a different profile, with bins concentrated in the med-high ME. The total phase space displacement induced by bisulfite, quantified as Kolmogorov-Smirnov Distance (KS D) = 0.60 between the two full phase-space distributions, is an order of magnitude larger than the biological signals identified in the subsequent sections of this study.

Having established the phase-space framework and quantified the contribution of the bisulfite induced artifact in brain tissue, we next applied true-mC entropy analysis to a proof-of-concept biological model for which the genetic perturbation is well defined: primary clear cell renal cell carcinoma (ccRCC) paired with matched normal kidney tissue from the same individual. Figure 2a shows the two-dimensional ME-MML phase-space distributions computed under True-mC for the normal kidney and ccRCC tumor. A difference map (normal minus tumor) enables direct comparison of both the magnitude and direction of change from normal to tumor across phase space. Both distributions occupy the expected inverted-U phase space structure bounded by the entropy envelope, confirming that their overall methylation-state distributions were broadly preserved.

**Fig. 2:**
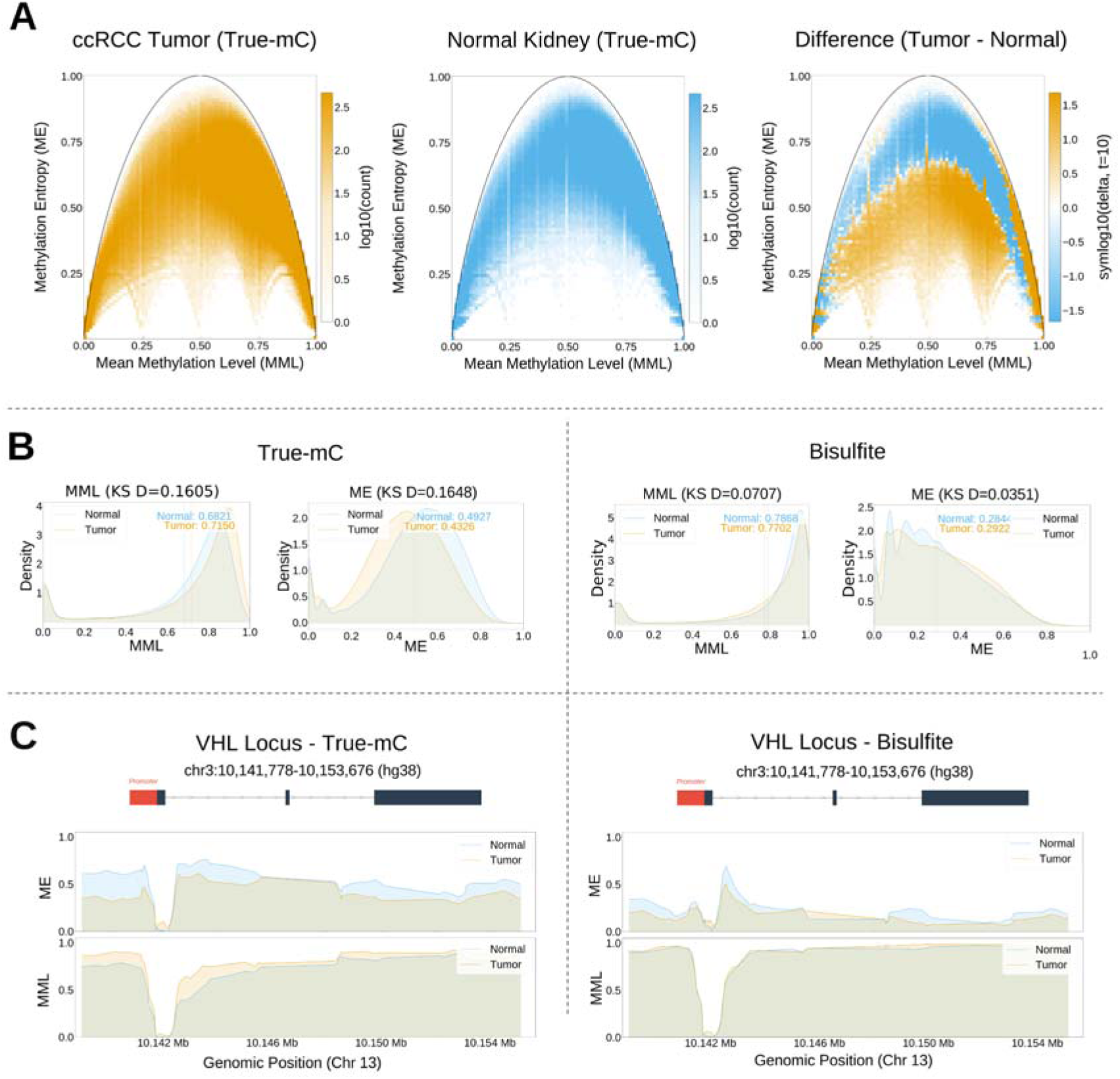
ccRCC versus normal kidney: phase-space redistribution and its erosion under bisulfite. **a** Top: 2D phase-space density (ME vs. MML, true-mC) for normal kidney (blue) and ccRCC tumor (orange). Right: signed log_₁₀_ difference map (normal minus tumor). **b** Distributions of MML and ME under true-mC (left) and bisulfite-like (right) for normal and tumor. **c** Locus-resolved MML and ME tracks at the remaining copy of the VHL gene body under true-mC (left) and bisulfite-like (right) for normal (blue) and tumor (orange). The ME differential visible under true-mC is absent under bisulfite.

The difference map may be viewed as a transition map from normal to tumor, with the excess bins in the normal tissue (blue) transitioning to the tumor excess bins (orange). The map displays three spatially distinct features. First, a concentrated orange feature runs along the right boundary of the arc at high MML (approximately 0.75-1.0), spanning a wide ME range. This represents a tumor-specific accumulation of bins at the highest methylation levels, consistent with focal hypermethylation gain in the tumor(58–60). This high-MML tumor-excess feature aligns with the widely reported tendency for cancers to acquire hypermethylation at specific loci(61, 62), as characterized across multiple tumor types(8). Second, normal-kidney-enriched bins (blue) are concentrated along the upper portion of the arc, spanning mostly intermediate-to-high MML values (roughly 0.3-0.75) at ME above approximately 0.65. This represents the high-entropy, moderately methylated loci characteristic of the normal kidney that the tumor has lost. Third, tumor-enriched bins (orange) form a broad band within the arc shifted towards lower ME values relative to normal, but with little shift in MML. The dominant redistribution towards a lower ME signal is inconsistent with stochastic erosion of the methylation landscape. Instead, it suggests coordinated, directional reorganization of methylation pattern fidelity.

Taken together, the difference map describes a bimodal DNA methylation redistribution in this tumor: a gain of highly methylated loci at the right arc boundary, and a simultaneous interior transition toward lower entropy at intermediate methylation levels, with depletion of the high-entropy normal-kidney states at the arc shoulder. We verified by simulation that the observed redistribution is not a result of random depletion in 5hmc(46, 47) and stochastic methylation in the kidney (Supplementary Fig. 8). In fact, our simulation shows that random redistribution increases tumor entropy, the opposite of what we observe in the real sample. Under bisulfite-like analysis, where 5hmC is interpreted as 5mC, this redistribution is eroded (Figure 2b). The tumor-normal separation visible in the True-mC ME distribution is compressed under bisulfite, while the MML distributions are comparatively less affected. This bisulfite-induced distortion is smaller in magnitude than in the brain (Figure 1c), consistent with the lower endogenous 5hmC of the kidney. However, it acts in the same direction, shifting bins toward the high-MML, low-ME corner of phase space. The difference in signal is directly observed in the remaining copy of VHL, the canonical ccRCC driver retained after 3p loss(46, 63, 64) (Figure 2c). At the VHL gene body, true-mC resolves a clear tumor-versus-normal ME differential with little accompanying change in MML. Under bisulfite-like analysis, this difference is largely erased, demonstrating at a single well-characterized driver locus how 5hmC conflation masks the structured entropy reorganization that true-mC recovers.

To decompose the ccRCC redistribution into its constituent phase-space movements, we defined two mutually exclusive bin sets based on the axis of primary displacement (Figure 3a). First, we defined bins that shifted in MML levels, filtered for substantial changes in MML (|dMML| > 50%), while controlling for ME (|dMME| < 10%). Second, we defined ME-shifted bins that showed substantial changes in methylation entropy (|dME| > 40%), while the MML was preserved (|dMML| < 10%). MML-shifted bins reflect classical methylation-level changes, while ME-shifted bins reflect changes in the variability across reads from different cells. The same average methylation level is maintained despite changes in the configurations of methylated and unmethylated states. Conventional differentially methylated regions (DMR)-based analytical frameworks would fail to identify these bins.

**Fig. 3:**
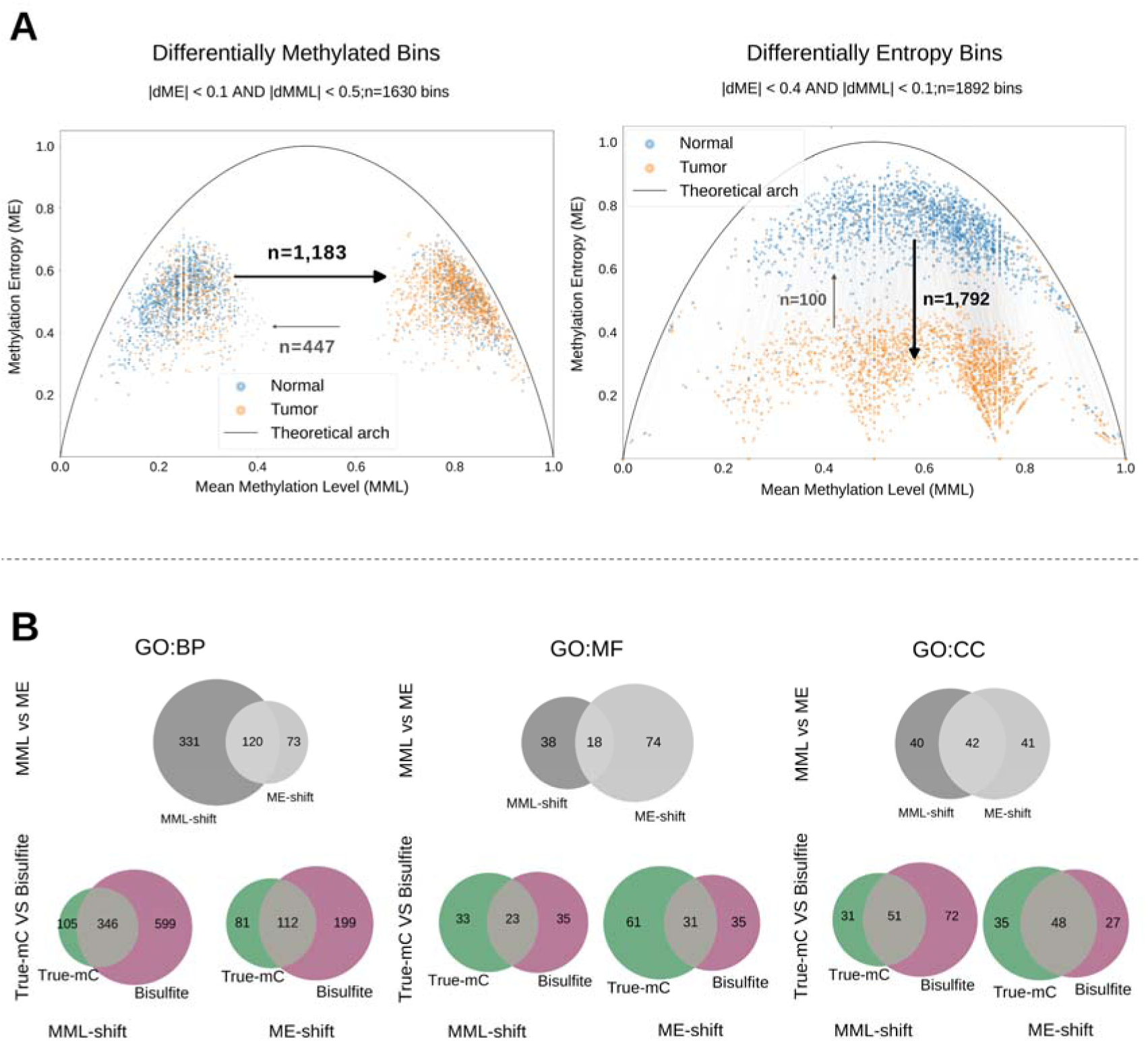
Axis decomposition of tumor-normal methylation changes and GO pathway recovery. **a** Phase-space scatter of MML-shifted bins (|ΔMML| > 0.5, |ΔME| < 0.1; n = 1,630) and ME-shifted bins (|ΔME| > 0.4, |ΔMML| < 0.1; n = 1,892), mutually exclusive by construction. MML-shifted: hypermethylation-to-hypomethylation ratio approximately 2.6:1. ME-shifted: entropy contraction-to-expansion ratio approximately 18:1. **b** Venn diagrams comparing genes recovered from MML-shift versus ME-shift bins across biological process (BP), molecular function (MF), and cellular component (CC) gene ontologies (GO). The lower Venn diagrams compare the overlapping GO terms recovered by the true-mC and bisulfite-like methods.

The directionality of these two sets differs markedly. Among MML-shifted bins, methylation gain in the tumor (n = 1,183) outnumbers loss (n = 447) by approximately 2.6-to-1. Among ME-shifted bins, entropy loss (n = 1,792) outnumbers entropy gain (n = 100) by approximately 18-to-1. This extreme directional shift serves as the empirical signature of a deterministic rewiring process. The tumor reorganizes its 5mC patterns into highly coordinated configurations without altering the average methylation level.

Gene ontology (GO) enrichment analyses computed for each bin set identified largely concordant pathway-level signals between true-mC and bisulfite (Figure 3b). The Venn diagrams show overlapping gene recovery between MML-shift and ME-shift sets. The molecular function ontology shows the lowest overlap, indicating that the two phase-space axes capture genuinely distinct molecular programs. Histone-modifying activity appears among top ME-shift molecular function terms, mechanistically consistent with a chromosome 3p loss model: SETD2 loss results in reduced H3K36me3 deposition. This histone modification serves to recruit DNMT3B to transcribed gene bodies, and its loss disrupts their stochastic methylation by.

Without this entropic contribution, intragenic methylation maintenance stabilizes certain patterns which manifest as entropy contraction (41, 59, 65, 66). The dot plots in Supplementary Figs. 9 and 10 confirm that bisulfite and true-mC enrichment values track closely across most top-ranking terms in all three ontologies. The identified pathways are largely the same across measurement schemes, but the phase-space architecture is systematically mislocalized.

We next evaluated our framework in a high-5hmC environment where bisulfite conflation is most acute. We investigated psilocybin-induced neuroplasticity in the mouse mPFC. Unlike the fixed genetic drivers of ccRCC, psilocybin acts as a transient pharmacological stimulus, inducing widespread synaptic remodeling that can persist for weeks after the compound has been metabolized(43). To capture the stabilization of these epigenetic changes, we analyzed samples 72 hours following a single intraperitoneal administration of psilocybin, long after the pharmacological effect had decayed. As anticipated for a dynamic pharmacological response, the magnitude of epigenetic redistribution was much smaller than in the stability-driven ccRCC samples, requiring the high fidelity of True-mC.

The phase-space difference map for psilocybin versus control is shown in Figure 4a with only a small fraction of bins (∼0.1%) showing distinct modulation. Although the redistribution is modest, psilocybin-enriched bins preferentially occupy regions of higher methylation entropy along the upper boundary of the phase-space arc, whereas control-enriched bins are concentrated at lower methylation entropy within the arc interior. This pattern suggests a directional shift toward higher methylation entropy after psilocybin exposure, consistent with the enhanced plasticity associated with psychedelics(43).

**Fig. 4:**
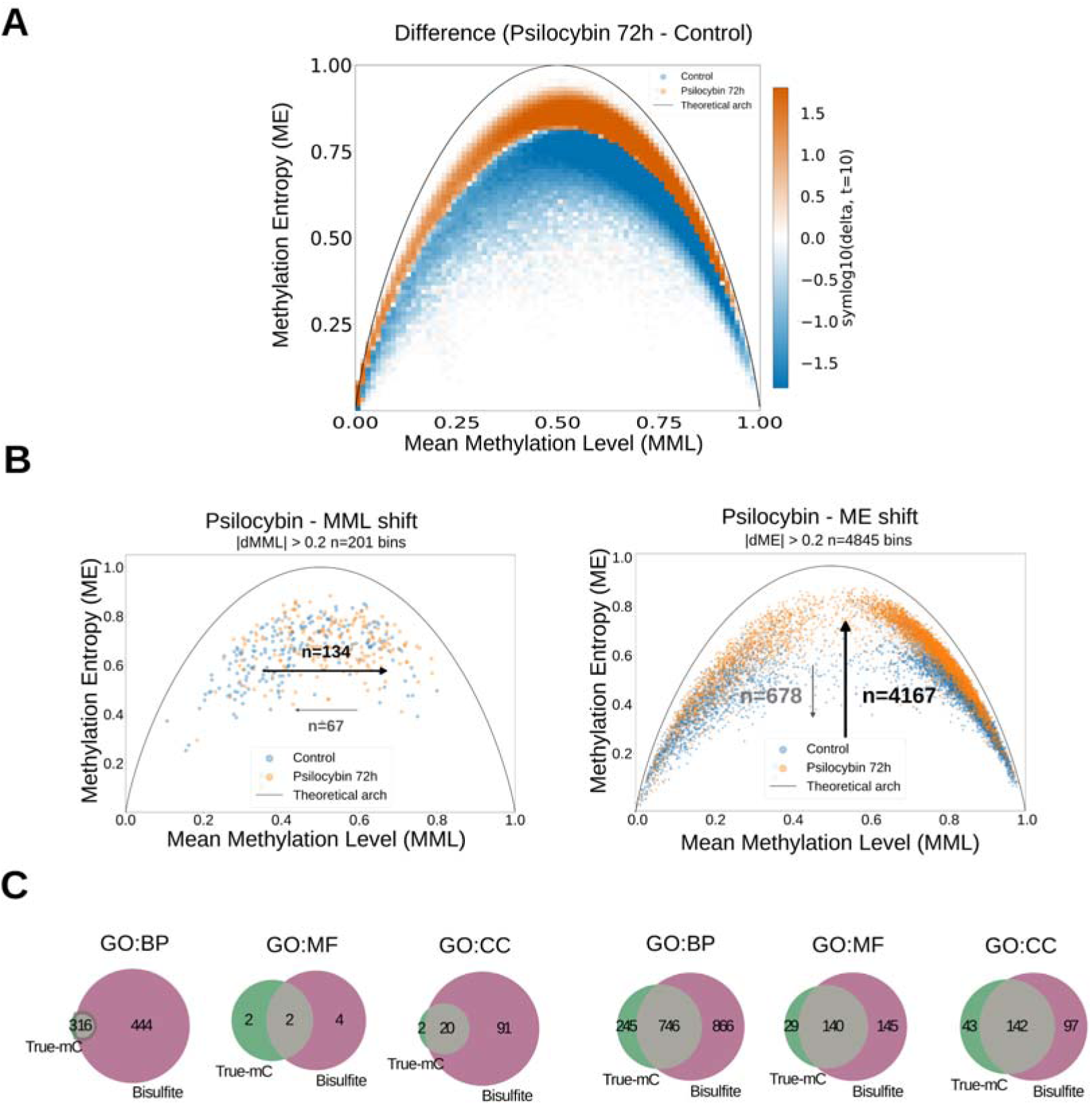
Psilocybin-induced entropy changes in prefrontal cortex and bisulfite misclassification. **a.** Signed log_₁₀_ difference map (psilocybin minus control) in phase space (ME vs. MML, True-mC, 4-CpG bins). Psilocybin-effect excess (orange) traces the upper arc; control excess (blue) accumulates in the interior. **b.** Phase-space scatter of MML-shifted bins (|ΔMML| > 20%; n = 201) and ME-shifted bins (|ΔME| > 20%; n = 4845). **c.** Venn diagrams comparing genes identified under true-mC (green) versus bisulfite-like (purple) for MML-shifted bins (left) and ME-shifted bins (right) across BP, MF, and CC ontologies.

Filtered bin analysis more clearly resolves the directionality of the psilocybin-associated methylation changes (Figure 4b). Because the effect size was smaller than in the ccRCC comparison, lower filtering thresholds were applied. The MML-shifted and ME-shifted bin sets were not defined as mutually exclusive, so bins showing coordinated changes along both axes could contribute to both analyses. Among bins that changed by more than 20% ME, entropy gain in psilocybin-treated animals (n = 4167) outnumbers entropy loss (n = 678) by approximately 6-to-1. Among bins with changes of more than 20% in MML, gains (n = 134) outnumber losses (n = 67) by approximately 2 to 1. The contrast in directional consistency is mechanistically important: psilocybin-induced neuroplasticity is associated with a more directionally coherent signal along the ME axis than along the MML axis, the reverse of the typical relationship in homeostatic tissue maintenance. At well-methylated loci, the predominant psilocybin-associated signal is an ME gain, indicating increased methylation-pattern heterogeneity without a comparable shift in mean methylation level. This is the empirical signature expected for active TET-mediated 5mC oxidation in progress, in which the structured 5mC architecture is transiently destabilized as part of the neuroplastic response(18, 67, 68).

GO enrichment from both bin sets is heavily neuronal and synaptic. MML-shifted bins recover homophilic cell-cell adhesion, neurogenesis, and neuronal development, with synapses, postsynaptic density, presynaptic, and asymmetric synapses at the cellular component level (Supplementary Fig. 11). ME-shifted bins recover neuron differentiation, neurogenesis, neuron development, and nervous system development at the biological process level, with synaptic membrane, postsynaptic membrane, microtubule, and cell projection at the cellular component level (Supplementary Fig. 12). The enrichment is biologically coherent with psilocybin-induced prefrontal neuroplasticity and is captured by both axes through partially distinct gene programs.

The contrast between true-mC and bisulfite-like gene recovery exposes a profound methodological vulnerability of bisulfite sequencing in 5hmC-dynamic biological processes (Figure 4c). In MML-dynamic bins, the bisulfite-like analysis identifies substantially more GO terms than true-mC. Although GO term counts can be inflated by ontology redundancy and by region-to-gene mapping, the scale of the discrepancy is striking. At the biological process level, 444 terms are unique to the bisulfite-like analysis, only 16 are shared, and only a handful are unique to true-mC. This imbalance is also evident across the MF and CC ontologies. In ME-dynamic bins, the same pattern persists but is proportionally less extreme: 866 terms are unique to the bisulfite-like analysis, 746 are shared, and 245 are unique to true-mC.

Consistent with the ternary concordance in Fig. 1b (true-mC r = 0.94 vs bisulfite r = 0.71), the terms unique to bisulfite are likely dominated by 5hmC conflation rather than true 5mC signal. This is expected mechanistically: when 5hmC is deposited in a coordinated manner, collapsing it into the 5mC pool alters inter-read pattern structure while leaving mean methylation comparatively preserved, so entropy is the more corrupted axis. The phase-space distributions (Figs. 1c, 2b), in which the ME axis separates under true-mC but compresses under bisulfite while MML is comparatively stable, indicate that in 5hmC-dynamic tissues the entropy readout of bisulfite is the most vulnerable to contamination.

## DISCUSSION

This study quantitatively assesses how the inability of bisulfite sequencing to distinguish 5mC from 5hmC biases measurements of methylation level and methylation entropy in a context specific manner. We introduce ShannonPore, a generally applicable tool for true-mC based methylation phase-scape analysis from nanopore sequencing data. Although true-mC is not ground truth, it provides closer approximation to the fully resolved three-state model than bisulfite (r = 0.94 vs 0.71). Direct comparison of bisulfite-like and true-mC MML/ME landscapes in mouse brain tissue demonstrated that bisulfite substantially suppressed ME while inflating MML. This distortion was not random measurement noise. TET deposits 5hmC at consistent regions-but at variable CpG sites across reads. Collapsing this structured 5hmC signal with the 5mC signal creates apparent inter-read uniformity, reducing entropy while increasing apparent methylation level. Bisulfite conflation, therefore, introduces a directional, tissue-dependent bias into entropy estimates rather than a symmetric error.

The ccRCC analysis illustrates the value of resolving true 5mC architecture in a low-5hmC, genetically driven system. In the matched tumor-normal pair, true-mC entropy analysis revealed a marked contraction of the tumor methylation landscape, with ME-shifted loci accounting for over half of the differential bins and an approximately 18:1 imbalance toward lower-entropy states. Because this observation derives from a single deeply characterized ccRCC pair, we present it as a proof-of-concept rather than a cohort-level generalization. Accordingly, the mechanistic model proposed below should be viewed as a candidate mechanism to be tested in larger cohorts across molecular subtypes and clinical stages.

The ccRCC entropy contraction is consistent with a model in which 3p chromosomal loss and consequent SETD2/PBRM1 haploinsufficiency drive structured maintenance reorganization rather than stochastic erosion of the methylation landscape. In this model, loss of SETD2 depletes H3K36me3 across gene bodies, withdrawing the chromatin signal that recruits DNMT3B to transcribed intragenic regions. Removing this stochastic *de novo* component leaves DNMT1 to maintain existing patterns with high fidelity without adding new pattern variability. The result is entropy contraction at preserved mean methylation: coordinated patterns replace heterogeneous ones because the principal source of stochastic variation is selectively eliminated(41, 59).

Our general observation for the ccRCC sample is that modulation in ME carries mechanistic information, and that the tissue acquires a more rigid methylome in the tumor. These observations were recently supported by independent evidence across substantially larger cohorts. Rossi et al. quantified local epipolymorphism across 207 tumor and normal kidney samples using bisulfite sequencing and provided independent cohort-scale evidence that methylation pattern disorder carries functional information in ccRCC(69). Although their results cannot confirm the direction of a true-mC entropy change, they show that methylation-pattern disorder carries functional, regulatory information in ccRCC beyond the methylation level. In another recent study, Hackett et al.(70) analyzed methylation entropy at the protocadherin (PCDH) gene cluster across 122 ccRCC tumors and 63 adjacent normals from 68 individuals. They report a significant reduction in entropy in tumors despite using orthogonal entropy metrics. This possibly distinguishes ccRCC from high-entropy malignancies such as lung and colorectal cancer(8, 61), refining the prevailing view that malignancy inherently increases methylation entropy.

The psilocybin-treated brain provides the complementary case: a high-5hmC, dynamically remodeled tissue state in which bisulfite and true-mC interpretations diverge sharply. Unlike the fixed genetic perturbation in ccRCC, psilocybin acts as a transient pharmacological stimulus associated with neuroplastic remodeling. In this context, psilocybin increased ME in the mPFC, with entropy gains greatly outnumbering losses among ME-modulated loci. The gain bias was stronger along the ME axis than along the MML axis, indicating that methylation entropy was the more sensitive readout of the underlying response. At well-methylated loci, the predominant psilocybin-associated signal was ME gain, indicating increased methylation-pattern heterogeneity without a comparable shift in mean methylation level. This pattern is consistent with active TET-mediated 5mC oxidation in progress, in which the structured 5mC architecture is transiently destabilized as part of the neuroplastic response(20, 71, 72). The extent of bisulfite discordance in the psilocybin dataset was substantial. Bisulfite conflation is blind to coordinated oxidation of methylated regions, missing the resulting methylation-pattern disorder. This effect was much less pronounced in ccRCC, where endogenous 5hmC is lower and pathway-level agreement between bisulfite and true-mC was stronger.

Taken together, the ccRCC and psilocybin analyses indicate that the methodological cost of bisulfite sequencing for entropy analysis is not a fixed offset. It scales with the endogenous 5hmC content and dynamics of the tissue under study. In low-5hmC contexts such as kidney, bisulfite-derived analyses may preserve pathway-level biological relevance even when they misattribute part of the entropy signal. In 5hmC-rich and 5hmC-dynamic contexts such as brain, the same measurement strategy can substantially alter both the magnitude and interpretation of methylation entropy. This distinction has implications beyond the two systems studied here. Entropy-based methylation analyses in embryonic development, aging, neurodegeneration, and other cytosine-remodeling states may be similarly blind to active demethylation dynamics. By separating true 5mC architecture from 5hmC-dependent distortion, true-mC resolved entropy analysis provides a more precise framework for interpreting epigenetic heterogeneity across biological contexts.

## ACKNOWLEDGEMENTS

We thank Sapir Margalit and Ofri Ranen for their help in the early steps of this project.

## AUTHOR CONTRIBUTIONS

U.B. performed data analysis, developed the computational tool, generated sequencing data, and contributed to manuscript writing. E.K., N.G., J.J., and A.G. contributed to data analysis. A.G. and J.D. contributed to sequencing data generation. S.V. contributed to the development of the computational tool. A.S., G.U.N., and B.L. contributed to animal handling and psilocybin administration. Y.R. and Y.E. conceived and supervised the study and edited the manuscript. All authors reviewed and approved the manuscript.

## SUPPLEMENTARY DATA

Supplementary Data are available at NAR online.

## DATA AVAILABILITY

Raw nanopore sequencing data for the ccRCC dataset have been deposited in the NCBI Sequence Read Archive under accession number PRJNA1196660. Processed methylation entropy data for the ccRCC dataset, together with raw and processed data for the psilocybin mouse dataset, have been deposited in NCBI’s Gene Expression Omnibus under accession number <u>GSE342266</u>.

## CODE AVAILABILITY

Methylation entropy and mean methylation level were computed using ShannonPore, an open-source platform developed for this study and available at https://github.com/ebensteinLab/ShannonPore. All analysis scripts used to generate the figures in this paper are available at <u>DOI:10.5281/zenodo.21835446</u>. Any inquiry about the tool or script should be sent to.

