## Supplementary information file for "Beyond Bisulfite Sequencing: Resolving 5-hmC with Nanopore Sequencing Unmasks the True Methylation Entropy Landscape"

**AUTHOR INFORMATION**

Uri Bertocchi[1], Eyal Katz[2], Jonathan Jeffet[2], Assaf Grunwald[3], Neiv Gabay[2], Jasline Deek[3], Sujal Verma[3], Amit Shwartz[5,6], Gali Umschweif–Nevo[5], Bernard Lerer[6], Yael Roichman[2,3,*], Yuval Ebenstein[1,3,4*]

[1]Sagol School of Neuroscience, Tel Aviv University, Tel Aviv-Yafo, Israel

[2]Raymond & Beverly Sackler School of Physics & Astronomy, Faculty of Exact Sciences, Tel Aviv University, Tel Aviv, Israel

[3]Raymond & Beverly Sackler School of Chemistry, Faculty of Exact Sciences, Tel Aviv University, Tel Aviv, Israel

[4]School of Biomedical Engineering, The Iby and Aladar Fleischman Faculty of Engineering, Tel Aviv University
[5]School of Pharmacy, Faculty of Medicine, The Hebrew University of Jerusalem, Jerusalem, Israel

[6]Hadassah BrainLabs Center for Psychedelic Research, Hadassah Medical Center, Hebrew University, Jerusalem, Israel

### SUPPLEMENTARY NOTES

#### Supplementary Note 1: Calculating mean methylation level (MML) and methylation entropy (ME)

To analyze epigenetic variability within a specific genomic bin containing a defined number of CpG sites, we use two metrics: Mean Methylation Level (MML) and Methylation Entropy (ME). While MML measures the baseline methylation abundance, ME captures the underlying spatial complexity and pattern heterogeneity of the reads.

The standard approach to evaluating a genomic region is calculating its Mean Methylation Level (MML), which measures the average proportion of methylated cytosines across all sequenced reads within a given bin. It is defined as:

$$MML = \frac{\Sigma_{reads}\Sigma_{CpG\ sites} methylation\ level}{\Sigma_{reads} site\ number}$$

However, MML, as a standalone metric, has a significant limitation: it is a bulk average that collapses distinct cellular subpopulations. As illustrated in Figure 1a, many distinct molecular methylation patterns can yield the same cumulative MML value, obscuring critical epigenetic variations.

To overcome this limitation and resolve the underlying pattern configurations, we introduce another metric - Methylation Entropy (ME). ME is the Shannon entropy of the empirical distribution of methylation patterns, normalized by the number of CpG sites in the bin. It is defined as:

$$\text{ME} = \frac{-1}{N_{CpG}}\sum_{i=1}^{2^{N_{CpG}}} P_i log_2 P_i$$

Where $P_i$ is the empirical probability for the i-th methylation pattern in the current bin, $N_{CpG}$ is the number of CpG sites within the bin, and $2^{N_{CpG}}$ is the total number of possible methylation patterns in a bin. In the case of ternary ME, as presented in Figure S7, this parameter becomes $3^{N_{CpG}}$ due to the 3 possible states at each site: C, 5mC, and 5hmC.

To associate methylation entropy with sample properties, it is important to map it along the genome, as is commonly done for methylation levels. By dividing the genome into bins and evaluating both metrics across all bins, we can characterize the sample's behavior in the ME-MML phase space. This framework provides a pattern-level information-theoretic measure that complements the mean methylation level and can reveal differences in read-level configurational diversity not captured by MML alone.

### SUPPLEMENTARY FIGURES

#### Comparing ShannonPore to existing nanopore methylation entropy tools

Current nanopore-based tools for assessing methylation heterogeneity, such as modkit and PoreMeth2, were not designed to isolate the 5hmC contribution and do not implement the Shannon-entropy framework used here. To establish how ShannonPore's output relates to these tools, we computed ME and MML for the same genomic bins using all three tools and directly compared the resulting values, together with their respective MML-ME phase-space distributions, relative to the theoretical Shannon entropy envelope. All three tools' parameters were set to 4 CpGs per bin with a minimum coverage of 16x. 5hmC modifications were counted as unmodified cytosines. Consequently, Modkit and PoreMeth2, which used a sliding-window approach across the genome, produced correlated but non-identical methylation estimates (ME) and methylation likelihood (MML). Additionally, the phase-space points generated by Modkit and PoreMeth2 occasionally fell outside the theoretical arch. Their entropy outputs are therefore tool-specific heterogeneity metrics rather than the normalized Shannon entropy of the read-level methylation-pattern distribution. Two consequences follow. First, sliding-window definitions place a variable number of CpGs in each window, so the state space over which entropy is normalized is not fixed and the resulting values are not comparable across bins of differing CpG density. Second, both tools returned points outside the theoretical entropy envelope, which is not attainable by any valid normalized Shannon entropy and indicates that the reported quantity is not bounded by the entropy of the underlying pattern distribution. ShannonPore's constant-CpG binning enforces a fixed state space per bin and therefore keeps all estimates inside the envelope.

**Supplementary Fig. 1: Comparison of ShannonPore, modkit, and PoreMeth2 estimates of methylation entropy and methylation level.**

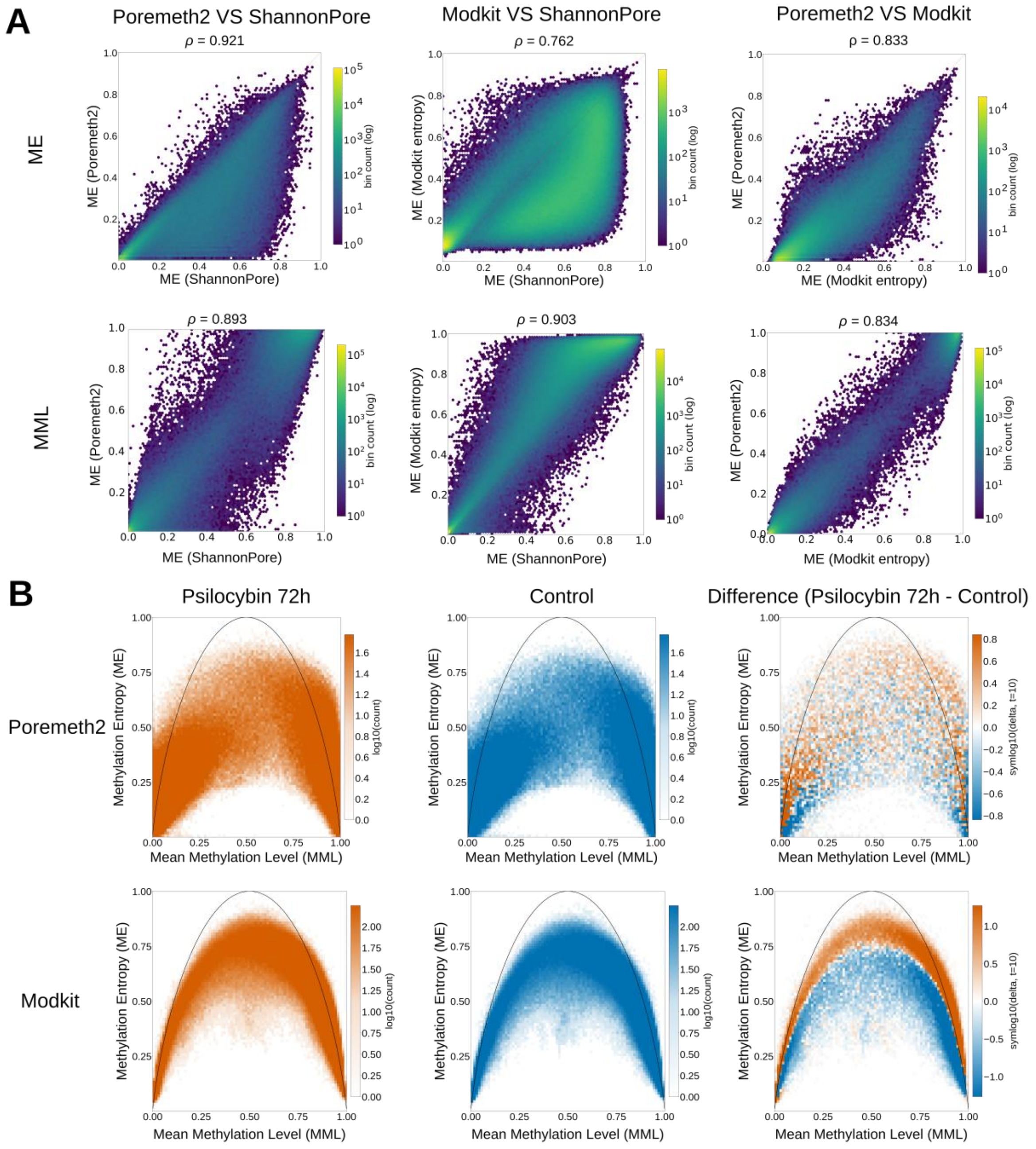


**a** Pairwise comparisons of ME (top row) and MML (bottom row) calculated by ShannonPore, modkit, and PoreMeth2 for matched genomic bins, shown as 2D density heatmaps (log-scaled bin counts), with Spearman's $\rho$ indicated above each panel. **b** MML–ME phase-space plots for control and psilocybin samples calculated using modkit and PoreMeth2. The black curve marks the theoretical upper bound on normalized binary Shannon entropy for a given MML.

**Optimization of genomic bin size for methylation entropy estimation**

To accurately capture the informational landscape of DNA methylation patterns, the choice of genomic binning strategy must balance regional CpG density with physical coverage depth. As shown in Fig S2 Panel A, using a fixed genomic window size (e.g., 150 bp) introduces a severe estimation error in regions with high CpG density; when coverage is constrained to 30x; splitting a dense cluster into a constant-size bin artifactually inflates the state space relative to the available reads, causing a steep drop-off in reliable information extraction. Conversely, adopting a "Constant CpG" strategy stabilizes the estimation error across varying densities by maintaining a controlled, uniform state-space size $2^N$ per bin. However, truncating the context window to a finite number of $N$ sites risks systematic information loss due to the omission of long-range correlations. As quantified in Panel B, this excess entropy drops exponentially and effectively vanishes when the window size matches or exceeds the intrinsic correlation length of 6 sites, as derived from the MML correlation decay in Panel C. Thus, optimizing the binning architecture requires a "Constant CpG" approach to bound the estimation error under fixed coverage, while ideally setting the number of sites N > 6 to completely mitigate maximum entropy (ME) digitization loss.

**Supplementary Fig. 2: Considerations in choosing bin size**

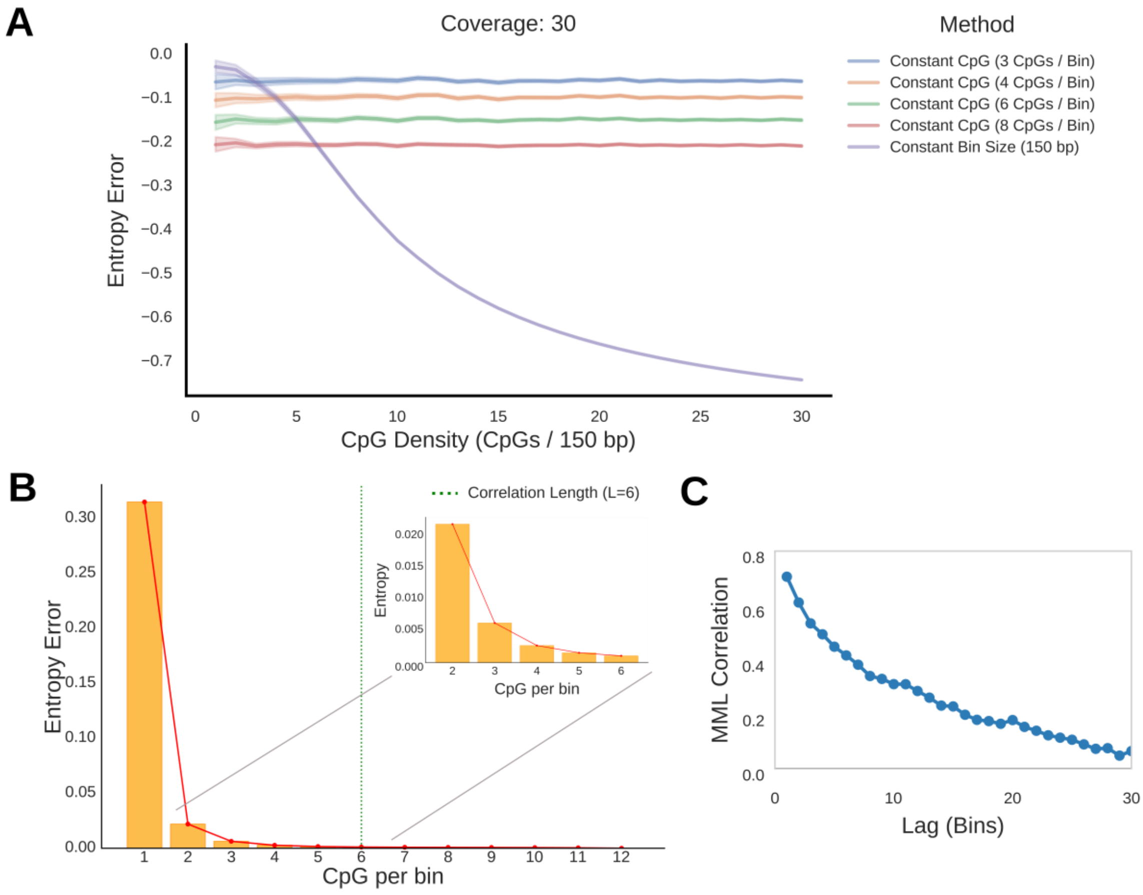


**a** ME estimation error vs CpG density. Four lines shows constant number of CpG per bin and one line shows a constant number of base pairs per bin. All in coverage 30x . **b** Entropy loss vs CpG sites per bin for autocorrelation length = 6 sites. **c** Autocorrelation function plot for bin methylation values across the genome. Each point represents the distance between two bins and the average correlation for this given distance.

The choice of bin size involves a trade-off between two sources of error. Sampling bias increases as the bin size increases, because fewer reads span the full bin. Information loss increases as the bin size decreases, because the spatial structure between sites is no longer captured. To find the bin size that minimizes the combined error, we plotted both error terms and their sum as a function of bin size, and marked the bin size that yields the lowest total error. We show here two scenarios: correlation lengths of 6 and 10 bins, both with coverage of 30x. In both cases, under the present assumptions and 30x coverage, the total modeled error is minimized near 3 CpGs per bin.

**Supplementary Fig. 3: Assessing the error related to bin size and coverage.**

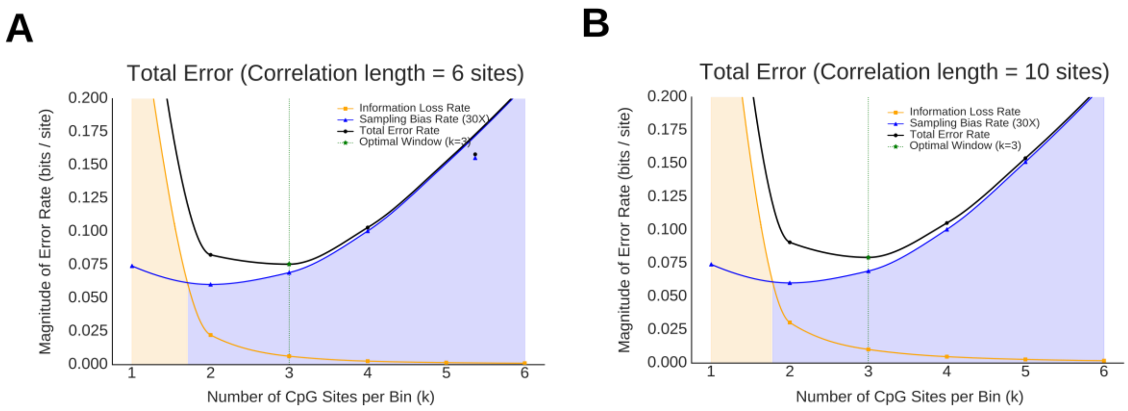


Each panel presents the magnitude of ME error as a function of the number of CpG sites per bin. Sampling bias error (blue line) and information-loss error (yellow line) are summed (black line) to show the total error. The shaded areas show which error rate is larger and are colored accordingly. **a** Correlation length 6 sites, 30x coverage: optimum at bin size 3. **b** Correlation length 10, 30x coverage: optimum at bin size 3.

However, this analysis does not account for two important considerations. The first is that our long reads allow us to capture long-range correlations, and the second is the richer accessible entropy space accessible by doubling the number of possible methylation configurations in 4 vs. 3 CpG bins. Overall, the price in added estimation error when moving to 4 CpG bins is on the order of 3%, and in our view, merits the use of 4 CpG bins. Support for this choice comes from a quantitative, information-capacity-based comparison of the two bin sizes. Under finite-coverage constraints, such as 30x depth, the empirical Shannon entropy is systematically underestimated. This sampling bias accounts for 8% of the total error  for 3-site bins and 11% for 4-site bins (Figure S4). The true theoretical information capacity of a bin is given by its size in bits, meaning 3 bits for a 3-site bin and 4 bits for a 4-site bin. Factoring in the structural bias, the net usable entropy is defined as the maximum capacity minus absolute systematic sampling penalty: $H_{usable} \;=\; k \times (1 - \Delta H_{bias})$ For the 3-site configuration, this yields 2.76 bits of true resolvable information capacity. Conversely, the 4-site configuration, despite facing a slightly higher percentage penalty due to its larger state-space meaning larger sampling bias, preserves 3.56 bits of usable information capacity. Computing the ratio of these two physical entropy bounds reveals a 1.29-times net increase in total accessible information capacity. This quantitative gain shows that expanding the bin size to 4 sites structurally overrides the nominal increase in sampling noise, maximizing the conditional information rate per site.

**. Supplementary Fig. 4: Information Capacity comparison of 3 sites and 4 sites bins**

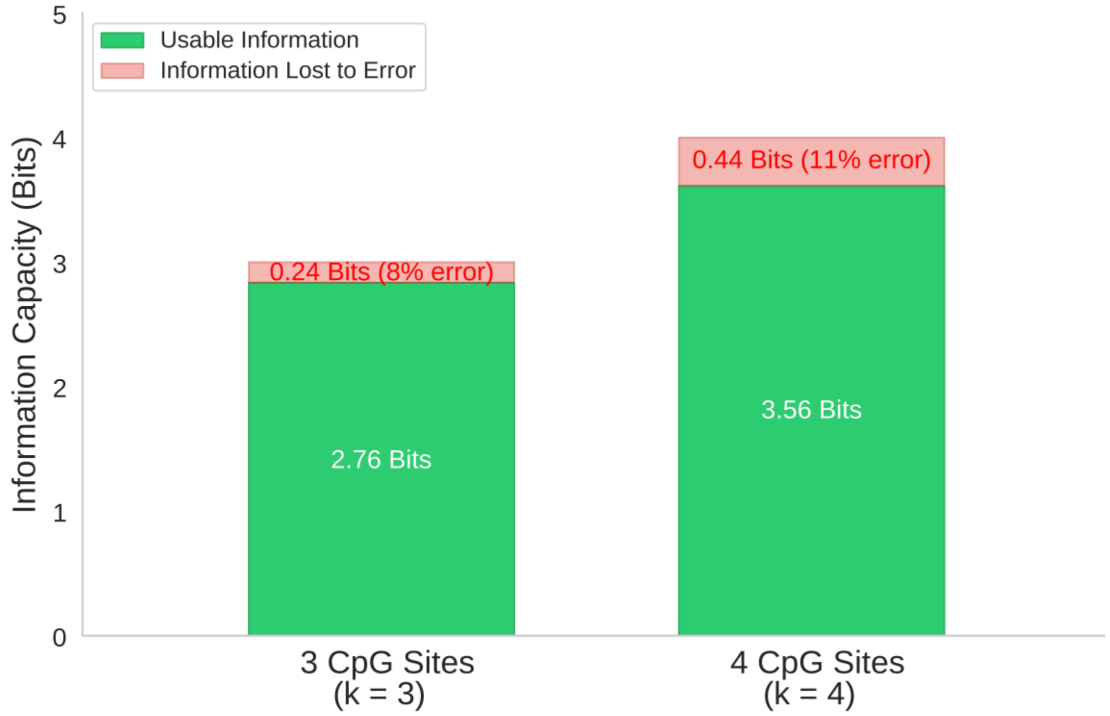


Stacked bars show the theoretical bit capacity of each bin size (3 bits for 3-CpG bins, 4 bits for 4-CpG bins) partitioned into usable information (green) and information lost to finite-coverage sampling bias (red) at 30x coverage.

This work uses nanopore sequencing with reads exceeding several Kbp. However, applying our strategy to short-read sequencing (~150 bp) may pose challenges in certain genomic regions with sparse CpG distribution. A constant-CpG binning strategy requires that the CpGs assigned to a bin co-occur within a single sequencing read. At low CpG densities, individual reads may not span the entire bin, leading to data loss. To assess this, we examined the genome-wide distribution of inter-CpG spacing (hg38) and the physical size, in base pairs, of 4-CpG bins, and compared these to a 150-bp read-length reference. The majority of 4-CpG bins fall within 150 bp, and 33% of all CpG sites occur at local densities exceeding 4 sites per 150-bp window, indicating that constant-CpG bins are mostly compatible with short-read sequencing and that the long read lengths achieved in this study result in no substantial loss of informative CpGs.

**Supplementary Fig. 5: Bin size histogram and CpG sites number histogram of hg38 reference**

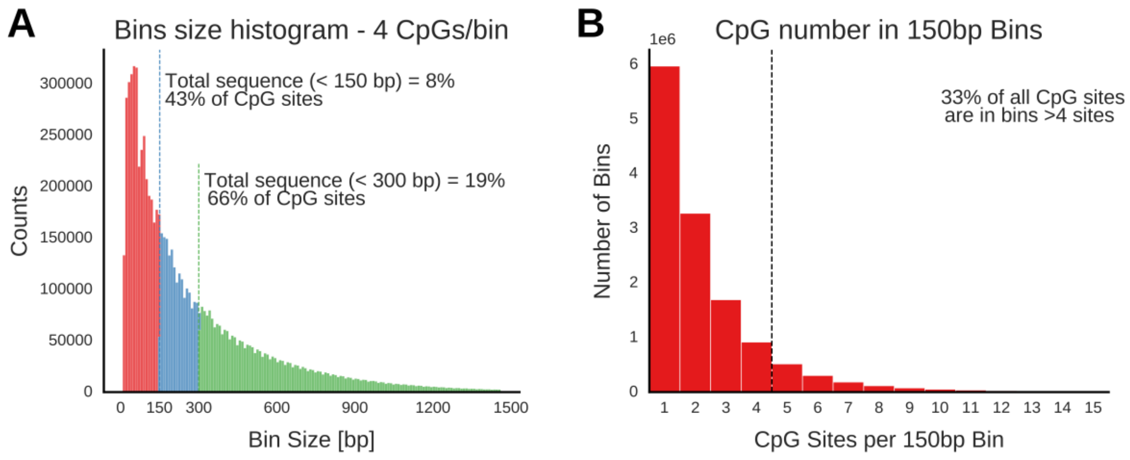


**a** Histogram of 4-CpG bin sizes (bp) across the genome, colored by span: 0–150 bp (red), 150–300 bp (blue), and >300 bp (green). **b** Histogram of CpG counts within 150-bp genomic windows.

### Global epigenetic composition in the studied samples

Differences in phase-space distributions between conditions could, in principle, reflect a global shift in the relative abundance of C, 5mC, and 5hmC, rather than locus-specific reorganization of methylation patterns. To rule this out, we quantified the genome-wide fraction of CpG calls assigned to each of the three states for every sample in both datasets. Saline- and psilocybin-treated mice showed near-identical global CpG-state composition, indicating that gross genome-wide compositional changes are limited in this dataset; however, this does not exclude cell-type composition shifts or regional redistribution of modifications, as described for Figure

4. In contrast, the ccRCC tumor showed reduced global 5hmC and increased 5mC relative to the normal kidney, consistent with previously reported 5hmC loss in kidney cancer[1]. A simulation for random reassignment of CpG states to recapitulate the global transition form normal to tumor is shown in Figure S8, confirming that the phase space redistribution observed in our study is not random.

**Supplementary Fig. 6: Global cytosine modification composition across samples.**

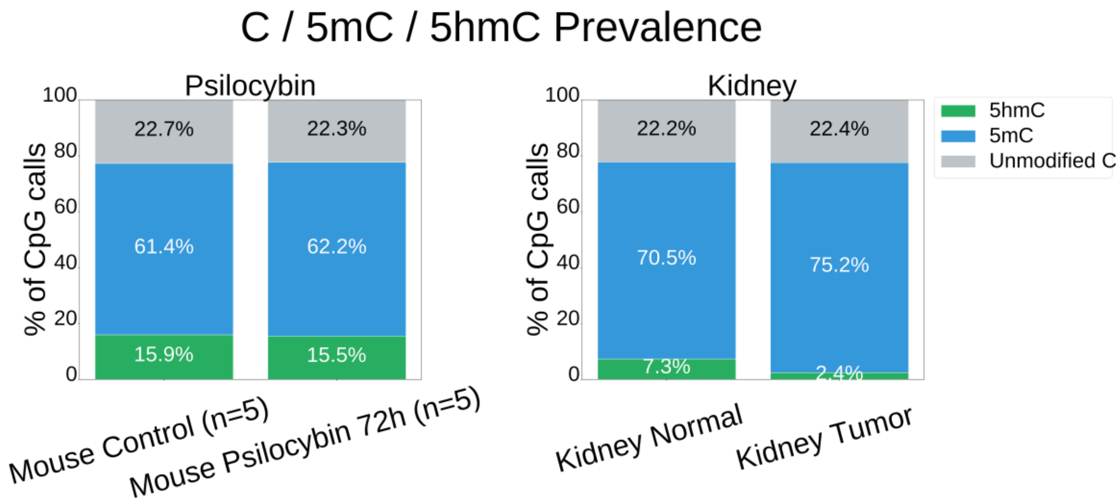


Bar plots showing the genome-wide fraction of CpG calls assigned to C, 5mC, and 5hmC for each sample in the psilocybin cohort (control and psilocybin-treated) and the ccRCC cohort (normal kidney and tumor).

#### *Adding 5hmC to the model*

To systematically decouple the structural organization of the modified methylome, this framework models the multidimensional interplay between the mean methylation level (MML) and the mean hydroxymethylation level (MHL). ME is still computed from binary C/5mC patterns; the ternary plot (Figure S7) is a visualization of how this entropy varies as a function of MML and mean hydroxymethylation level.  This approach captures the distribution of configurational disorder without altering the underlying binary state space used to compute the entropy. The theoretical boundary of this space is defined by a mathematical ceiling (red dashed line) representing an idealized, highly synchronized system in which genomic regions transition between purely unmodified and fully modified states, bypassing highly disordered intermediate states. Comparing the control group vs. the psilocybin brain group, and normal kidney tissues vs. ccRCC tissues, reveals distinct landscape shifts following both psilocybin administration and oncogenic transformation. These shifts are consistent with altered relationships among methylation, hydroxymethylation, and pattern-level methylation entropy.

**Supplementary Fig. 7: ME landscape in MML vs. MHL phase space.**

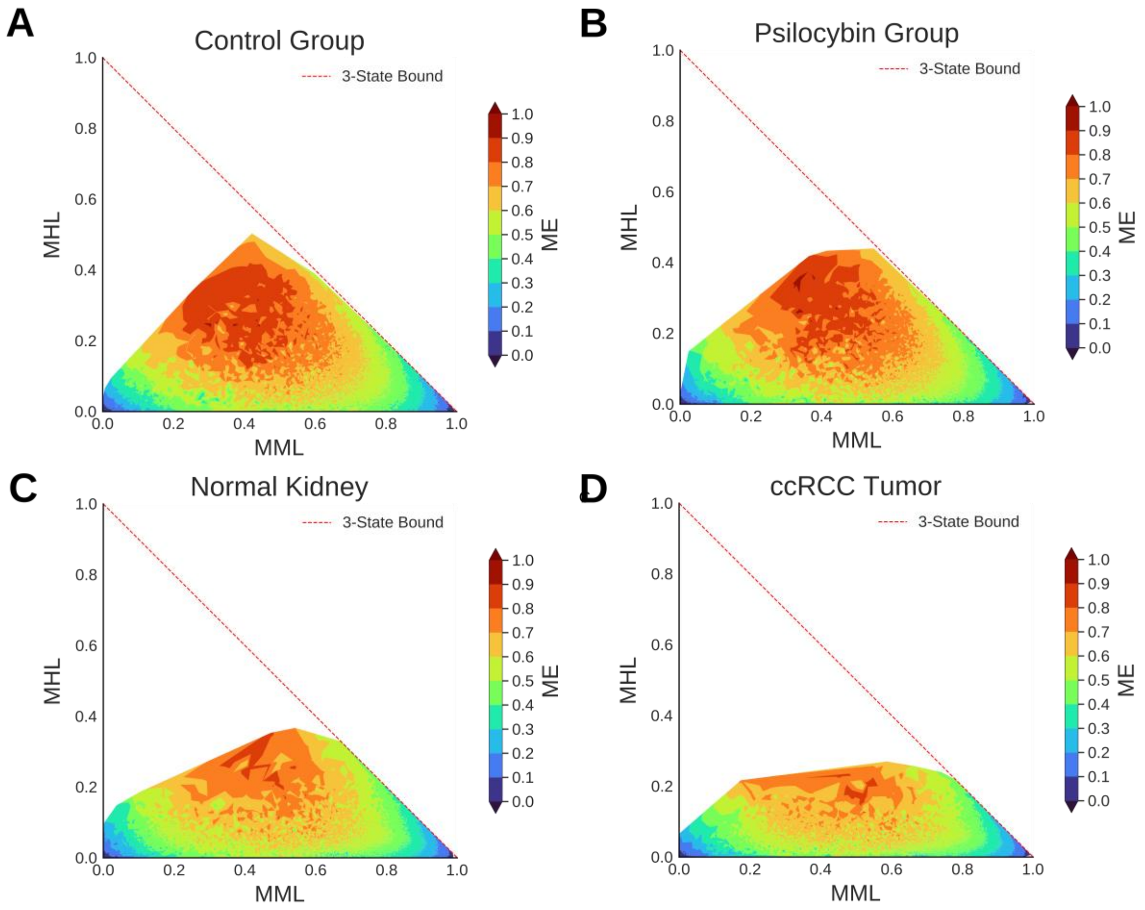


Bivariate density landscapes mapping 5mC (MML, x-axis) against mean 5hmC levels (MHL, y-axis). Methylation entropy (ME) is projected as a continuous surface across the coordinate space, with the color scale representing ME values from low (blue) to high (dark red). The red dashed line denotes the theoretical 3-State Bound, which defines the mathematical limits of the configuration space between these distinct epigenetic states. The top row displays the psilocybin experiment, comparing the **a** Control Group and **b** Psilocybin Admission group. The bottom row displays the kidney cancer (ccRCC) experiment, comparing **c** normal kidney tissue and **d** ccRCC tumor tissue.

**Simulated tumor: ruling out passive 5hmC depletion and stochastic re-methylation as drivers of the ccRCC entropy contraction**

The normal-kidney sample carries a higher global 5hmC level than the ccRCC tumor (Figure S6), and under the True-mC framework, loss of 5hmC reassigns CpGs from the modified to the unmodified-C state. We therefore tested whether the ME contraction observed in the tumor (Figures 2–3) could arise simply from passive, non-directed 5hmC depletion and stochastic re-methylation during tumor proliferation, rather than from enzymatically directed reorganization of the 5mC pattern.

To this end, we constructed a randomized control from the normal-kidney read-level state matrix. We randomly selected reads with a 5hmC call and converted them to unmodified C until the sample's global 5hmC level matched the tumor level, then randomly converted C calls to 5mC until the global 5mC level also matched the tumor level. This produced a "randomized normal" sample with the tumor's global cytosine composition (5hmC, 5mC, and C fractions matched within rounding) but with both the demethylation and re-methylation steps applied at random rather than at directed loci. We then compared the phase-space distribution of this randomized control with that of the real tumor.

The randomized normal reproduced the tumor's global modification levels but not its phase-space signature. On the MML axis, the three samples were nearly indistinguishable (KS D = 0.076 randomized-vs-tumor), as expected once global methylation levels are matched. However, the difference map (simulated tumor minus normal) lacks the directional, low-entropy interior consolidation characteristic of real tumor redistribution (Figure 2) and shows high ME excess along the arc in the tumor. Taken together, these results show that matching global 5hmC and 5mC levels by random reassignment is insufficient to reproduce the tumor’s phase-space structure, suggesting that locus-specific constraints beyond simple passive dilution are involved.

**Supplementary Fig. 8: Randomized 5hmC-depletion control rules out passive dilution as the source of ccRCC entropy contraction.**

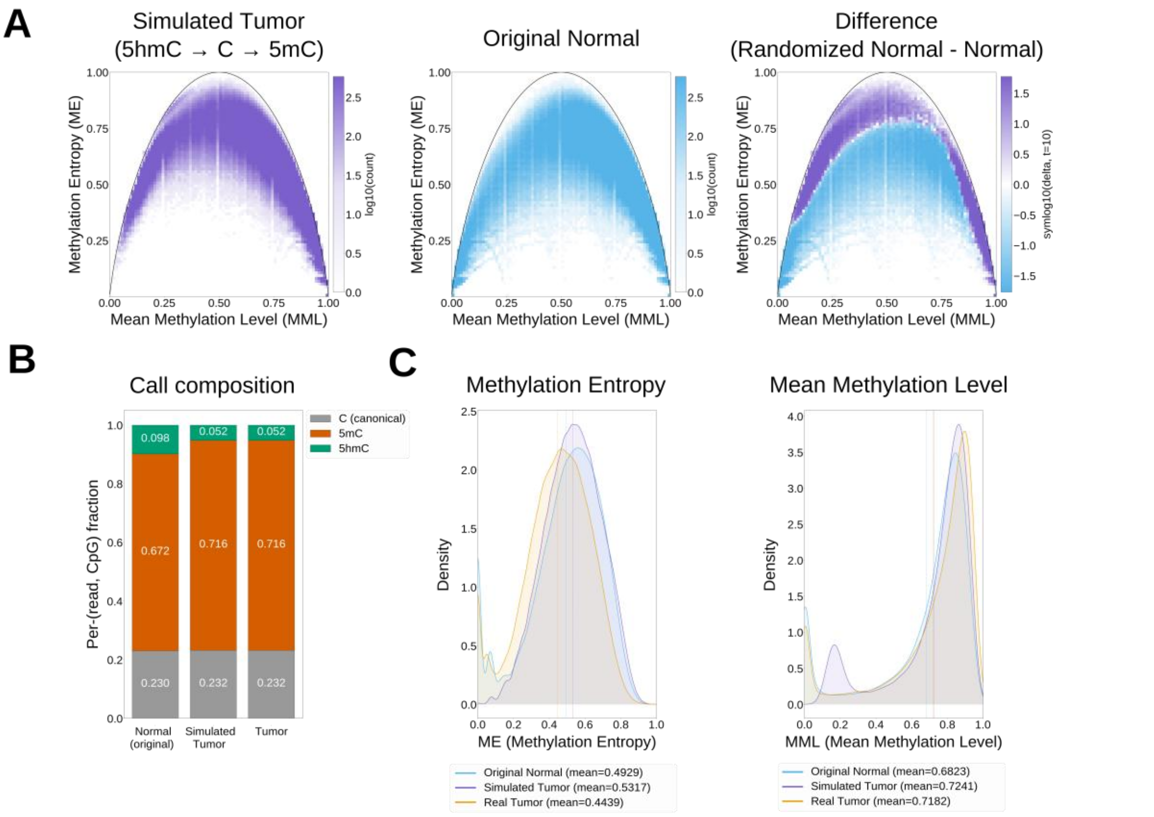


**a** True-mC phase-space density maps (ME vs. MML, 4-CpG bins) for the simulated (randomized) tumor (left, purple), the real normal kidney (middle, blue), and their signed $\log_{10}$ difference (right; simulated tumor minus normal). **b** Per-read CpG call composition (fraction of C, 5mC, and 5hmC) for the original normal kidney, the randomized normal, and the real tumor; global composition is matched between the randomized normal (simulated tumor) and the tumor. **c** Density distributions of ME (left) and MML (right) for the original normal (blue), randomized normal (orange/red), and real tumor (yellow), with per-distribution means indicated. The KS D statistic between the randomized normal and the real tumor is shown above each panel.

### Gene ontology enrichment concordance between true-mC and bisulfite-like for ccRCC MML-shift and ME-shift bins

To assess whether the choice of measurement framework affects the biological pathways recovered from the ccRCC MML-shift and ME-shift bin sets (Figure 3a), beyond the gene-level overlap already shown in the Venn diagrams (Figure 3b), we compared the enrichment significance of top-ranking GO terms under true-mC and pseudo-bisulfite directly. For each bin set and ontology (BP, MF, CC), we plotted the q-value under both frameworks for the top-ranking terms, sized by the number of genes contributing to each term. Across nearly all top-ranking terms in all three ontologies, true-mC and bisulfite-like q-values track closely, indicating that, unlike the phase-space architecture (Figure 2), the top pathway categories recovered here are broadly similar under both frameworks, although the underlying gene sets and term significance can differ. This concordance includes histone-modifying activity among the top ME-shift molecular function terms, consistent with the SETD2/H3K36me3/DNMT3B mechanism proposed in the discussion.

**Supplementary Fig. 9: Concordance of GO term enrichment between true-mC and bisulfite for ccRCC MML-shift bins.**

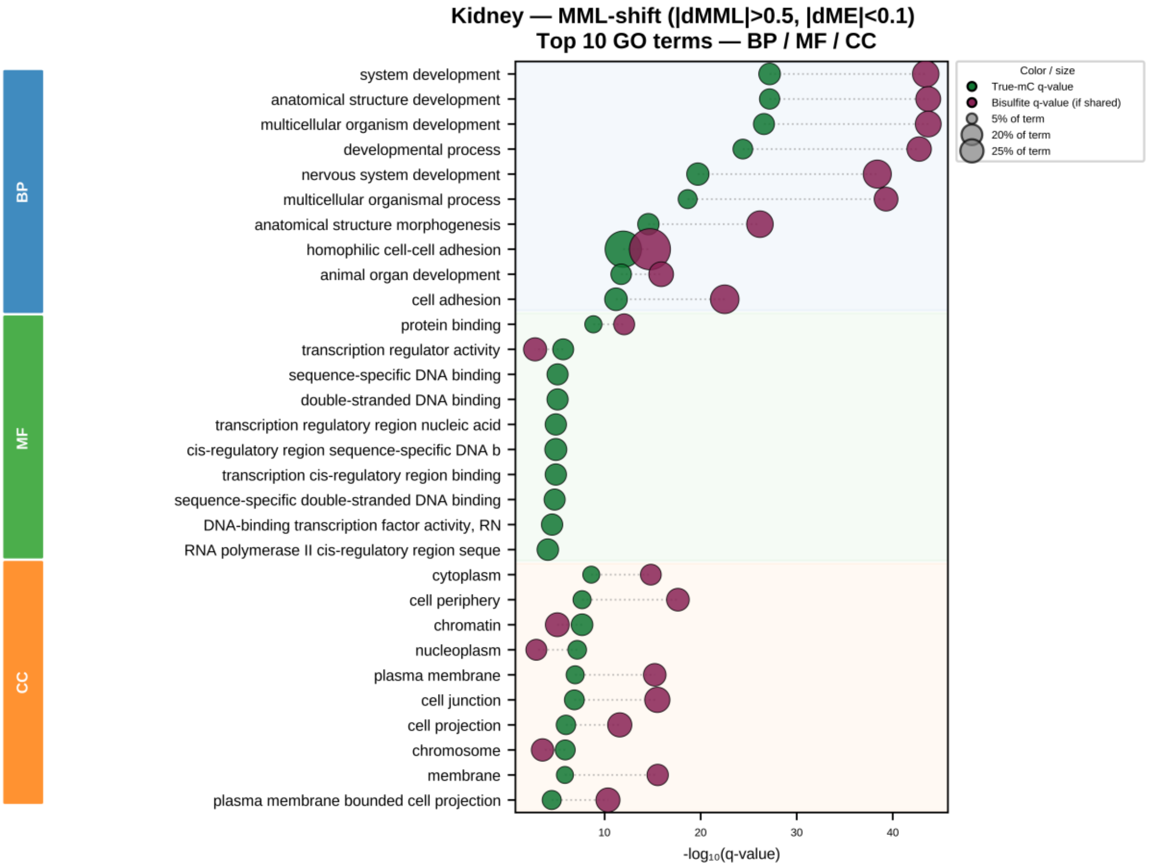


Dot plots of top-ranking enriched GO terms per ontology (BP, MF, CC; colored side bars) for a MML-shift bins (|ΔMML| > 50%, |ΔME| < 10%; n = 1,630). Blue dots show -$\log_{10}$(q-value) under True-mC; red dots show -$\log_{10}$(q-value) under pseudo-bisulfite for the same term. Dot size indicates the number of genes annotated to the term in the input bin set.

### Supplementary Fig. 10: Concordance of GO term enrichment between true-mC and bisulfite for ccRCC ME-shift bins

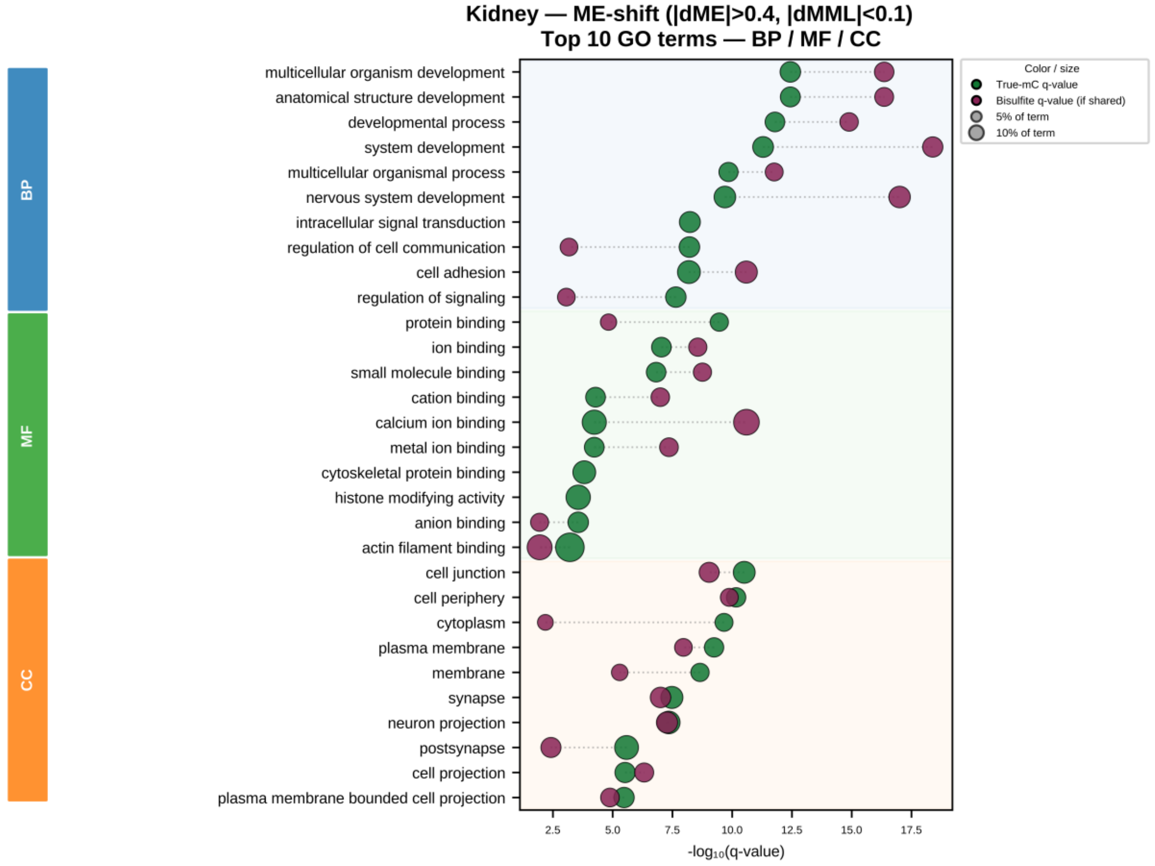


ME-shift bins (|ΔME| > 40%, |ΔMML| < 10%; n = 1,892). Blue dots show -$\log_{10}$(q-value) under true-mC; red dots show -$\log_{10}$(q-value) under pseudo-bisulfite for the same term. Dot size indicates the number of genes annotated to the term in the input bin set.

### Gene ontology enrichment of psilocybin-responsive ME-shift and MML-shift bins

To assess whether True-mC and pseudo-bisulfite recover the same biological pathways for the psilocybin-responsive bins described in the main text (Figure 4b), we performed GO enrichment analysis on both bin sets under each framework, across all three ontologies (biological process, BP; molecular function, MF; cellular component, CC). For each ontology, the top 10 enriched terms by true-mC q-value are shown, with the corresponding pseudo-bisulfite q-value plotted alongside wherever that term was also recovered under bisulfite.For MML-shift bins (Figure S10), top BP terms include homophilic cell-cell adhesion, neurogenesis, and neuronal development, with synapse, postsynaptic density, presynaptic, and asymmetric synapse terms recovered at the CC level.  For ME-shift bins (Figure S11), the top terms at the BP level are dominated by neuronal differentiation, neurogenesis, and nervous system development, and at the CC level by synaptic membrane, postsynaptic membrane, microtubule, and cell projection. Across both bin sets, enrichment is dominated by neuronal and synaptic biology, consistent with psilocybin-induced prefrontal neuroplasticity, though the magnitude of true-mC and bisulfite q-values frequently diverges even when the same term is recovered by both frameworks, reflecting partially distinct underlying gene sets. We note that the bisulfite-like analysis yielded a significantly higher number of enriched terms, most of which did not overlap with True-mC terms.

**Supplementary Fig. 11: Top GO terms for MML-shift bins (psilocybin vs. control).**

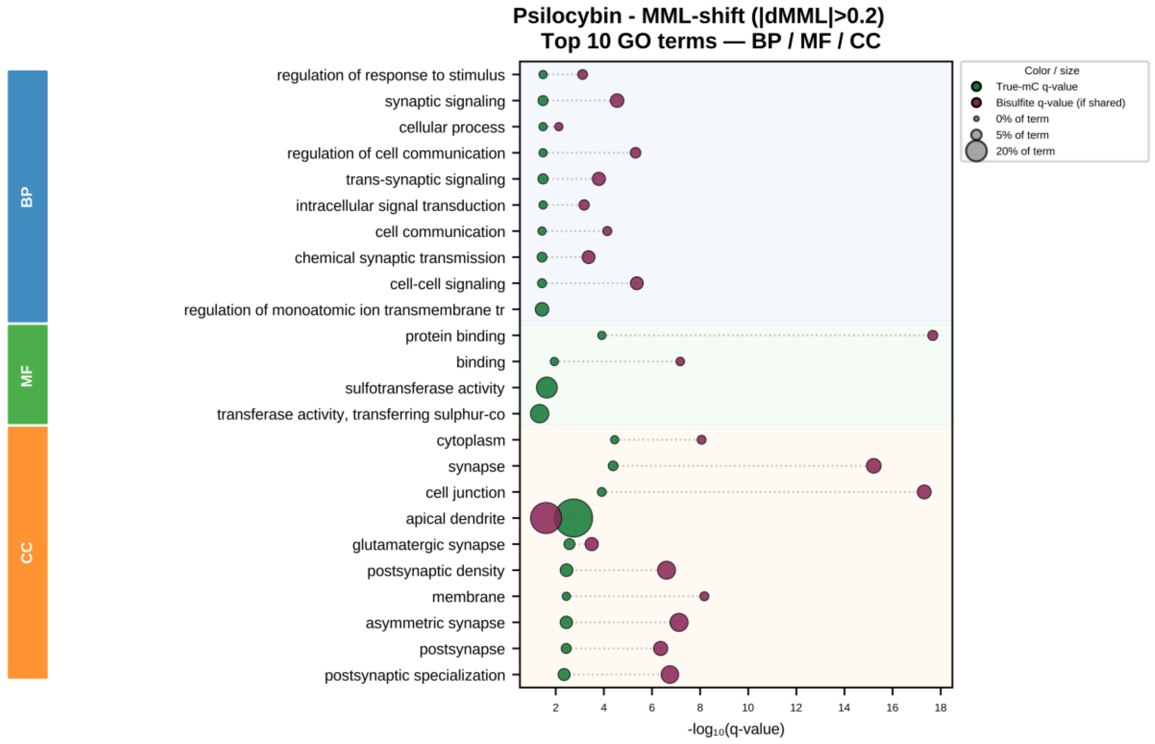


Dot plot of the top 10 enriched GO terms per ontology (BP, MF, CC; colored side bars) for bins with |ΔMML| > 20% (n = 201). Blue dots show -$\log_{10}$(q-value) under true-mC; red dots show -$\log_{10}$(q-value) under pseudo-bisulfite for terms that are also recovered in that framework. Dot size indicates the number of genes annotated to the term in the input bin set, as shown in the size legend.

**Supplementary Fig. 12: Top GO terms for ME-shift bins (psilocybin vs. control).**

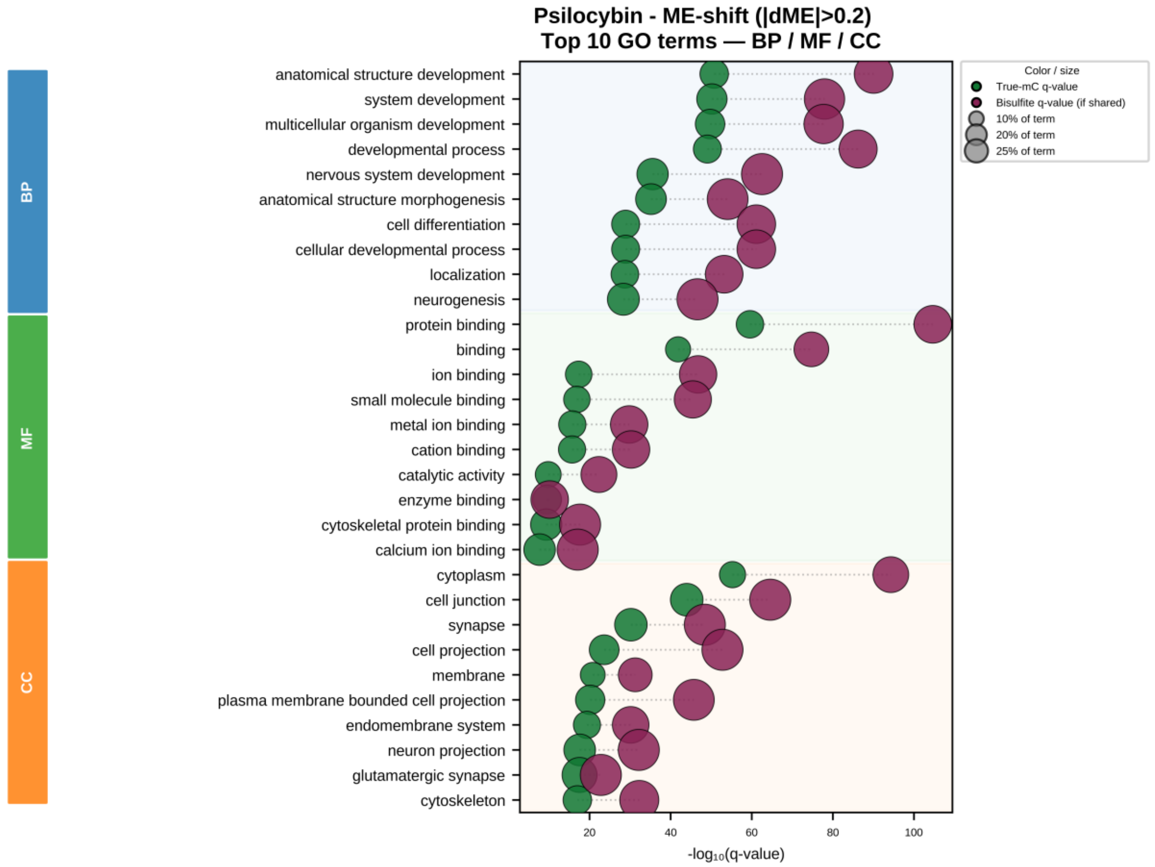


Dot plot of the top 10 enriched GO terms per ontology (BP, MF, CC; colored side bars) for bins with |ΔME| > 20% (n = 4845). Blue dots show -$\log_{10}$(q-value) under true-mC; red dots show -$\log_{10}$(q-value) under pseudo-bisulfite for terms that are also recovered in that framework. Dot size indicates the number of genes annotated to the term in the input bin set, as shown in the size legend.

#### Chromosome 3p loss in the ccRCC tumor sample

The SETD2-haploinsufficiency mechanism proposed to explain the entropy contraction observed in this tumor depends on this sample carrying a 3p deletion, since SETD2, PBRM1, BAP1, and VHL are all located on the short arm of chromosome 3. The genomic architecture of this tumor/normal pair was previously characterized by integrated Optical Genome Mapping and nanopore long-read sequencing, including genome-wide copy-number analysis in 500-kb bins, which identified a single-copy loss spanning the entire 3p arm in the tumor. Here we reproduce the relevant copy-number evidence from the nanopore dataset used in the present study to confirm the 3p deletion underlying our entropy analysis; these results were published by Margalit et al[1,2].

**Supplementary Fig. 13: Chromosome 3p copy-number loss in the ccRCC tumor.**

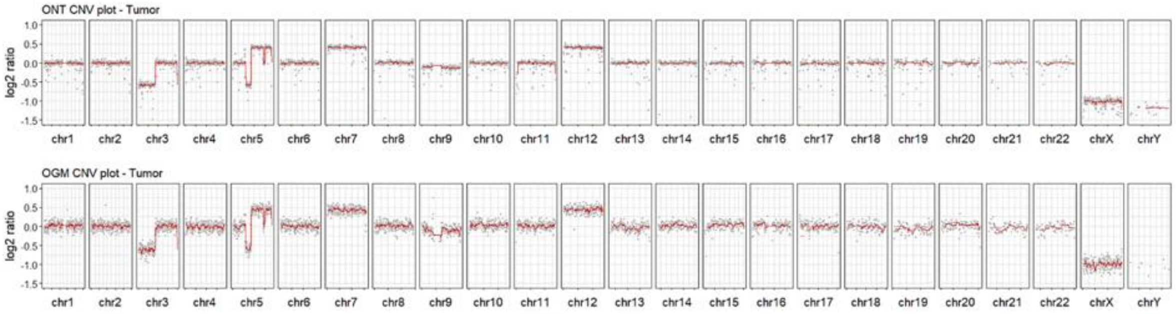


Comparative analysis of copy number variations (CNVs) in a ccRCC tumor, as detected by ONT and OGM. The plots show log2 of the copy ratio generated from ONT (top) and OGM (bottom) data. Data illustrate highly similar findings, pinpointing a significant DNA loss of the 3p arm of chromosome 3 and various losses and gains on chromosomes 5, 7 and 12.

### SUPPLEMENTARY TABLES

**Supplementary Table 1. Per-sample nanopore sequencing quality control metrics.** Total reads, fraction of primary reads aligned to the reference (% aligned), mean genome-wide coverage, and read length N50 computed from primary, non-supplementary alignments.

| Sample | Group | Total reads | % aligned | Mean cov | Read N50 (bp) |
|---|---|---|---|---|---|
| Sample-16 | control | 10,206,295 | 99.28% | 21.3× | 11,350 |
| Sample-18 | control | 7,742,295 | 87.82% | 11.8× | 8,932 |
| Sample-03 | control | 31,506,437 | 95.05% | 31.3× | 7,340 |
| Sample-04 | control | 14,844,744 | 94.09% | 16.9× | 12,536 |

| | | | | | |
|---|---|---|---|---|---|
| Sample-05 | control | 19,319,224 | 90.20% | 23.6× | 12,984 |
| Sample-06 | psilocybin_72h | 14,369,331 | 93.65% | 20.2× | 13,035 |
| Sample-07 | psilocybin_72h | 24,736,640 | 91.41% | 34.8× | 14,449 |
| Sample-08 | psilocybin_72h | 23,555,438 | 93.37% | 34.9× | 14,101 |
| Sample-09 | psilocybin_72h | 20,985,462 | 92.68% | 34.3× | 14,257 |
| Sample-10 | psilocybin_72h | 17,995,785 | 88.60% | 17.4× | 11,074 |
| ccRCC_normal | ccRCC | 17,242,669 | 98.07% | 20.3× | 15,287 |
| ccRCC_tumor | ccRCC | 32,051,007 | 97.09% | 36.3× | 18,503 |
